# Structure-optimized sgRNA selection with PlatinumCRISPr for efficient Cas9 generation of knock-outs

**DOI:** 10.1101/2022.12.15.520550

**Authors:** Irmgard U. Haussmann, Thomas C. Dix, David W. J. McQuarrie, Veronica Dezi, Abdullah I. Hans, Roland Arnold, Matthias Soller

**Affiliations:** School of Biosciences, College of Life and Environmental Sciences, University of Birmingham, Edgbaston, Birmingham, B15 2TT, United Kingdom; College of Life Science, Birmingham City University, Birmingham, B5 3TN, United Kingdom; Institute of Cancer and Genomics Sciences, College of Medical and Dental Sciences, University of Birmingham, Edgbaston, Birmingham, B15 2TT, United Kingdom; Birmingham Centre for Genome Biology, University of Birmingham, Edgbaston, Birmingham, B15 2TT, United Kingdom; Division of Molecular and Cellular Function, School of Biological Sciences, Faculty of Biology, Medicine and Health, University of Manchester, Michael Smith Building, Oxford Road, Manchester, M13 9PT, United Kingdom

**Keywords:** sgRNA/Cas9 optimization, secondary structure, retro-transposition

## Abstract

A single guide RNA (sgRNA) directs Cas9 nuclease for gene-specific scission of double-stranded DNA. High Cas9 activity is essential for efficient gene editing to generate gene deletions and gene replacements by homologous recombination. However, cleavage efficiency is below 50% for more than half of randomly selected sgRNA sequences in human cell culture screens or model organisms. We used in vitro assays to determine intrinsic molecular parameters for maximal sgRNA activity including correct folding of sgRNAs and Cas9 structural information. From comparison of over 10 data sets, we find major constraints in sgRNA design originating from defective secondary structure of the sgRNA, sequence context of the seed region, GC context and detrimental motifs, but we also find considerable variation among different prediction tools when applied to different data sets. To aid selection of efficient sgRNAs, we developed web-based PlatinumCRISPr, an sgRNA design tool to evaluate base-pairing and sequence composition parameters for optimal design of highly efficient sgRNAs for Cas9 genome editing named PlatinumCRISPr. We applied this tool to select sgRNAs to efficiently generate gene deletions in *Drosophila Ythdc1* and *Ythdf*, that bind to *N*^6^ methylated adenosines (m^6^A) in mRNA. However, we discovered, that generating small deletions with sgRNAs and Cas9 leads to ectopic reinsertion of the deleted DNA fragment elsewhere in the genome. These insertions can be removed by standard genetic recombination and chromosome exchange. These new insights into sgRNA design and the mechanisms of CRISPR-Cas9 genome editing advances efficient use of this technique for safer applications in humans.

## Introduction

Bacterially derived Clustered Regularly Interspaced Short Palindromic Repeats (CRISPR)-associated protein 9 (Cas9) from *Streptococcus pyogenes* provides a powerful tool for precise genome editing (Garcia-Doval and Jinek 2017; Jiang and Doudna 2017; Hille et al. 2018; Doudna 2020). To induce double stand-breaks in DNA at desired locations, the DNA scission enzyme Cas9 uses a guide RNA (gRNA) containing a 20 nucleotide complementary sequence to the genomic target site (protospacer), which also requires the protospacer adjacent motif (PAM) at the 3’end, comprised of an NGG sequence (whereby N is any nucleotide and G is guanine). In addition to the target complementary sequence (spacer) the gRNA also contains a constant crispr RNA (crRNA) sequence that base-pairs with *trans*-activating crRNA (tracrRNA). Alternatively, a single gRNA (sgRNA) can be used whereby the crRNA is fused to the tracrRNA through an artificial loop (Jinek et al. 2012; Cong et al. 2013).

High efficiency CRISPR-Cas9 mediated DNA scission is essential to generate mutants at high frequency in genetic screens and to provide the resource for efficient homologous recombination directed gene replacements. Large scale analysis of sgRNA efficiencies revealed the whole spectrum of on-target cleavage activities ranging from 0-100% arguing for a number of parameters that need to be correct for high efficiency cleavage leading to models incorporating weighing of features and/or thermodynamics of secondary structures (Hsu et al. 2013; Doench et al. 2014; Gagnon et al. 2014; Ren et al. 2014; Wang et al. 2014; Chari et al. 2015; Farboud and Meyer 2015; Hart et al. 2015; Housden et al. 2015; Moreno-Mateos et al. 2015; Varshney et al. 2015; Xu et al. 2015; Doench et al. 2016; Liu et al. 2016; Abadi et al. 2017; Gandhi et al. 2017; Chuai et al. 2018; Labuhn et al. 2018; Graf et al. 2019; Zhang et al. 2019; Michlits et al. 2020; Sledzinski et al. 2020; Trivedi et al. 2020; Xiang et al. 2021; Riesenberg et al. 2023). These studies identified that sequences with very low (≤35%) or very high (>80%) guanine-cytosine (GC) overall content were less effective indicating a critical aspect for binding energy in target scission. In addition, purines in the six nucleotides 5′ to the PAM substantially increased Cas9 cleavage efficiency, while pyrimidines and in particular uridine resulted in a lower efficiency (Ren et al. 2014; Wang et al. 2014; Housden et al. 2015; Graf et al. 2019). The lower efficiency of two uridine preceding the PAM site was further associated with premature termination of RNA Pol III (Graf et al. 2019), which can terminate after a stretch of four to six uridines (Gao et al. 2018). Moreover, changes in internal structure of sgRNA has been found associated with low activity (Moreno-Mateos et al. 2015; Thyme et al. 2016; Jensen et al. 2017). Recently, also bioinformatic approaches employing machine-learning have been used to improve prediction of sgRNA cleavage efficiencies (Xiang et al. 2021). These observations have helped to improve the design of sgRNAs to yield higher efficiencies and were incorporated into sgRNA design tools, but correlations between predictions and guide activity vary considerably (Haeussler et al. 2016; Labun et al. 2016; Sledzinski et al. 2020). Accordingly, available rules to predict sgRNAs are currently not sufficient to guarantee high cleavage efficiency and many sgRNA candidates scoring high fail to cleave efficiently (Haeussler et al. 2016; Labun et al. 2016; Labuhn et al. 2018; Sledzinski et al. 2020). In particular, the impact of sgRNA folding has not yet been analysed in detail and incorporated in web tools for sgRNA design. The X-ray crystal structure of Cas9 bound to sgRNA has been determined and indicates four regions of base-pairing termed tetraloop (tetraloop forms due to fusion of crRNA and tracrRNA) and stem loops 1-3 that might be important for its function (Anders et al. 2014; Nishimasu et al. 2014). This structure revealed many points of close interactions of the folded sgRNA with Cas9, but whether disruptions in the sgRNA structure would impact on Cas9 cleavage efficiency has not systematically been analyzed (Riesenberg et al. 2023). In particular, highly GC-rich guide RNAs could disrupt the rather weak secondary structure of the sgRNA bound by Cas9.

The CRISPR-Cas9 genome editing tool is widely used to generate knock-out mutants by introducing frameshifts. It has been recognized that introducing premature termination codons (PTCs) can induce use of alternative translation initiation sites (Tuladhar et al. 2019). In addition, in CRISPR-Cas9 engineered “knock-outs” of the m^6^A mRNA methyltransferase METTL3, it has been found that a functional ORF can be restored by altered splicing leaving considerable levels of m^6^A in mRNA (Poh et al. 2022). Likewise, compensatory responses have been observed involving upregulating genes and as a consequence causing stronger phenotypes than compared to removing a gene entirely (El-Brolosy et al. 2019; Ma et al. 2019). In addition, some genes have dual functions as protein and RNA (Hachet and Ephrussi 2004). Hence, introducing a frameshift will only remove the protein function. Likewise, many non-coding RNAs are present in introns suggesting that their expression is connected to the expression of the host gene (Boivin et al. 2018), but such relationships have not yet been explored comprehensively as they require more sophisticated genome editing (Deveson et al. 2017). Thus, for generating gene knock-outs, removing the transcription start site or the entire gene should be considered.

The sequence complementary guided targeting of the gRNA/Cas9 complex makes this an ideal tool for genome editing, but its use is currently limited by the low predictability to cut its target in the genome (Haeussler et al. 2016; Labuhn et al. 2018; Sledzinski et al. 2020). In particular, the role of RNA secondary structure of sgRNA has not been thoroughly evaluated and there is no sgRNA prediction tool that comprehensively takes RNA secondary structure of sgRNAs into account. To facilitate design of optimal sgRNAs, we developed an online tool incorporating all currently known parameters for sgRNA design including correct sgRNA folding (https://platinum-crispr.bham.ac.uk/predict.pl). Moreover, we provide guidelines to generate complete gene knockouts by generating deletions using dual sgRNAs. Taken together, these new sgRNA design and gene editing tools will help to develop CRISPR-Cas9 genome editing for safe application in humans.

## Results

### RNA secondary structure constraints limit sgRNA/Cas9 activity

Although sgRNA/Cas9 can cleave DNA efficiently, the first sgRNAs (L11GC and R13GC) we designed according to previously published guidelines did not cut the *pUC 3GLA Dscam 3-5* reporter we designed to study *Dscam* alternative splicing from introducing mutations by gap repair recombineering (**Fig. 1A,B, Supplemental Fig. S1A, Supplemental Table S1A**)(Hemani and Soller 2012; Haussmann et al. 2019). Similarly, the first sgRNAs flanking the *Drosophila Ythdf* gene, a reader for m^6^A mRNA methylation did not result in a deletion of the locus based on screening for loss of an RFP-marked transposon even though we had validated that the target sequence in this strain does not contain sequence polymorphisms (n=103, **Supplemental Fig. S2, Supplemental Table S1B**)(Balacco and Soller 2019). To determine the intrinsic molecular parameters for maximal sgRNA activity, we devised an in vitro assay to test the DNA scission efficiency of these two sgRNA based on in vitro transcribed sgRNAs and commercially available Cas9 using oligonucleotide and plasmid substrates containing matching protospacer sequences followed by a PAM.

**Figure 1:**
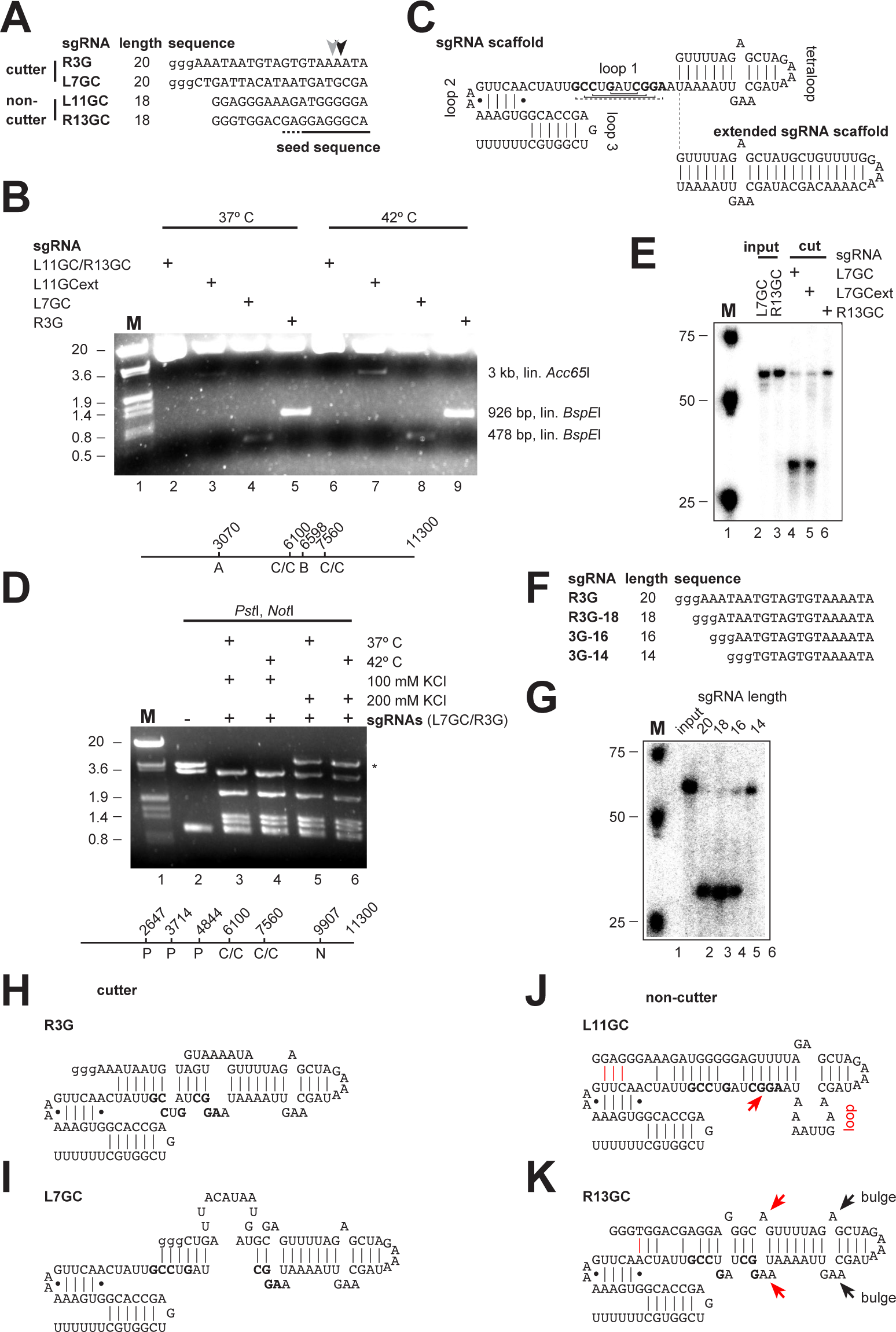
Sequence dependent in vitro cleavage of oligonucleotides and plasmid DNA by the sgRNA/Cas9 complex. (A) Sequences of sgRNAs with observed cleavage sites indicated by arrow heads. Small letter guanosines used for in vitro transcription are not present in the target DNA sequence. The seed sequence is indicated by a line at the bottom. (B) Agarose gel showing Cas9 mediated cleavage of the 11.3 kb Dscam 3-5 plasmid for 24 h with indicated sgRNAs. Plasmids were cut with either Acc65I (lanes 2, 3, 6 and 7) or BspEI (lanes 4, 5, 8 and 9) after Cas9 cleavage. The line at the bottom shows a map of the plasmid with restriction sites indicated. Size markers are EcoRI/HinDIII digested λ DNA of 20 kb, 3.6 kb, 1.9 kb and 0.8 kb. (C) Structure of the sgRNA scaffold from co-crystallization with Cas9 (Nishimasu et al., 2014). Vertical or horizontal lines indicate Watson-Crick base-pairing, and dots or dashed lines indicate non-Watson-Crick base-pairing. Nucleotides base-pairing in loop 1 are bold. Additional base-pairing found in the tracrRNA-crRNA heterodimer is indicated in the extended scaffold (Jinek et al., 2012). (D) Agarose gel showing Cas9 mediated cleavage of the 11.3 kb Dscam 3-5 plasmid for 24 h with indicated sgRNAs L7GC and R3G. Plasmids were cut with PstI and NotI. The star denotes incomplete cleavage by NotI and the line at the bottom shows a map of the plasmid with restriction sites indicated. Size markers are EcoRI/HinDIII digested λ DNA of 20 kb, 3.6 kb, 1.9 kb and 0.8 kb. (E) Denaturing acrylamide gel showing Cas9 mediated cleavage of synthetic oligonucleotides with indicated sgRNAs. (F) Sequences of sgRNAs with variable length. Small letter guanosines used for in vitro transcription are not present in the target DNA sequence. (G) Denaturing acrylamide gel showing Cas9 mediated cleavage for 1 h of synthetic oligonucleotides with indicated sgRNAs of variable length. (H-K) Structure of sgRNAs. Nucleotides base-pairing in loop 1 are bold. Red lines in J and K indicate potential base-pairing with nucleotides in loop 2. The red arrow in J indicates the sequence complementarity leading to a bulge in the tetraloop. The red arrows in K indicate a duplication of the bulge structure present in the tetraloop.

It has previously been shown that extending the tetraloop in the constitutive component of sgRNAs constituted by tracrRNA and crRNA enhances cleavage efficiency in vitro using oligonucleotide substrates (**Fig. 1C**)(Jinek et al. 2012). Introducing the extended sequence present in tracrRNA and crRNA into sgRNA (L7GCext) did not increase efficiency of cleaving a plasmid at 37°C, but increasing the temperature to 42°C enhanced cleavage by L7GCext (**Fig. 1B**). Increasing the salt concentration to 200 mM also did not result in enhanced cleavage by L7GC/R3G (**Fig. 1D**). In contrast, both L7GC and L7GCext could cleave an oligonucleotide, while R13GC did not (**Fig. 1E**). Using this assay also confirms the previous observation that sgRNAs of shorter length will lead to cleavage. In addition, three guanosines introduced at the 5’end of sgRNAs required for efficient in vitro transcription are tolerated (**Fig. 1F,G**)(Jinek et al. 2012).

The sgRNA scaffold adopts a typical fold when bound to Cas9 consisting of the bulged tetraloop, followed by small loop 1 and the more extended loops 2 and 3, which form a protective 3’end structure (**Fig. 1C**)(Nishimasu et al. 2014). The loop2/3 structure does not involve the uridines incorporated for termination of RNA Pol III driven expression from plasmids (**Fig. 1C**). When comparing the secondary structures of the four sgRNAs, we noticed that the two well-cutting sgRNAs L7GC or R3G maintained the secondary structure of the constitutive RNA part, while the non-cutting sgRNA L11GC disrupted the structure of the tetraloop (**Figs. 1H-K**). The effect of R13GC seems more subtle as it could cut in the oligonucleotide assay suggesting that the repeated bulge structure is the cause for its inefficiency, which is supported by X-ray crystal structure of the Cas9-sgRNA-DNA complex. Here, the bulge structure is recognized by Cas9 where Tyr_359_ base-stacks with G_43_, that also forms hydrogen bonds with Asp_364_ and Phe_351_, and Phe_351_ forms a hydrogen bond with A_42_ (Nishimasu et al. 2014). In addition, the sgRNAs initially used for deleting the *Ythdf* gene have a severely disrupted secondary structure (**Supplemental Fig. S2**).

When we systematically analysed genome sequences from *Drosophila* or humans for correct folding and activity of sgRNAs using the above parameters, about 50% of sgRNAs (241 from 481 and 503 from 973, respectively) did not fold properly. Also, only about 10-20 % of randomly selected sgRNAs exert high cleavage efficiency suggesting that correct folding could essentially contribute to high cleavage efficiency (Hsu et al. 2013; Doench et al. 2014; Gagnon et al. 2014; Ren et al. 2014; Wang et al. 2014; Chari et al. 2015; Farboud and Meyer 2015; Moreno-Mateos et al. 2015; Doench et al. 2016; Liu et al. 2016; Abadi et al. 2017; Chuai et al. 2018; Labuhn et al. 2018; Graf et al. 2019; Zhang et al. 2019; Michlits et al. 2020; Sledzinski et al. 2020).

When comparing the sequences of the cutting sgRNAs L7GC or R3G with the non-cutting L11GC and R13GC sgRNAs, we further noticed that L11GC and R13GC sgRNAs contained more guanosines, which in RNA can base-pair with C and U. To test if guanosines in the sgRNA limit Cas9 activity we increased their number in R3G to 13 to make sgRNA 13G (**Fig. 2A**). For the design of the R13 sgRNA, care was taken not to disrupt the tetraloop, but we noticed the potential to interfere with loop 2 (**Fig. 2B**, see below). We noticed, that sgRNA 13G is not capable of directing Cas9 cleavage if the target sequence is present in a 3 kb plasmid, but is active with a short oligonucleotide substrate (**Figs. 2A-D**). Likewise, adding a restriction enzyme together with sgRNA/Cas9 inhibited Cas9 in cleaving plasmid DNA suggesting that Cas9’s ability to scan DNA can be impaired separately from its ability to cleave DNA.

**Figure 2:**
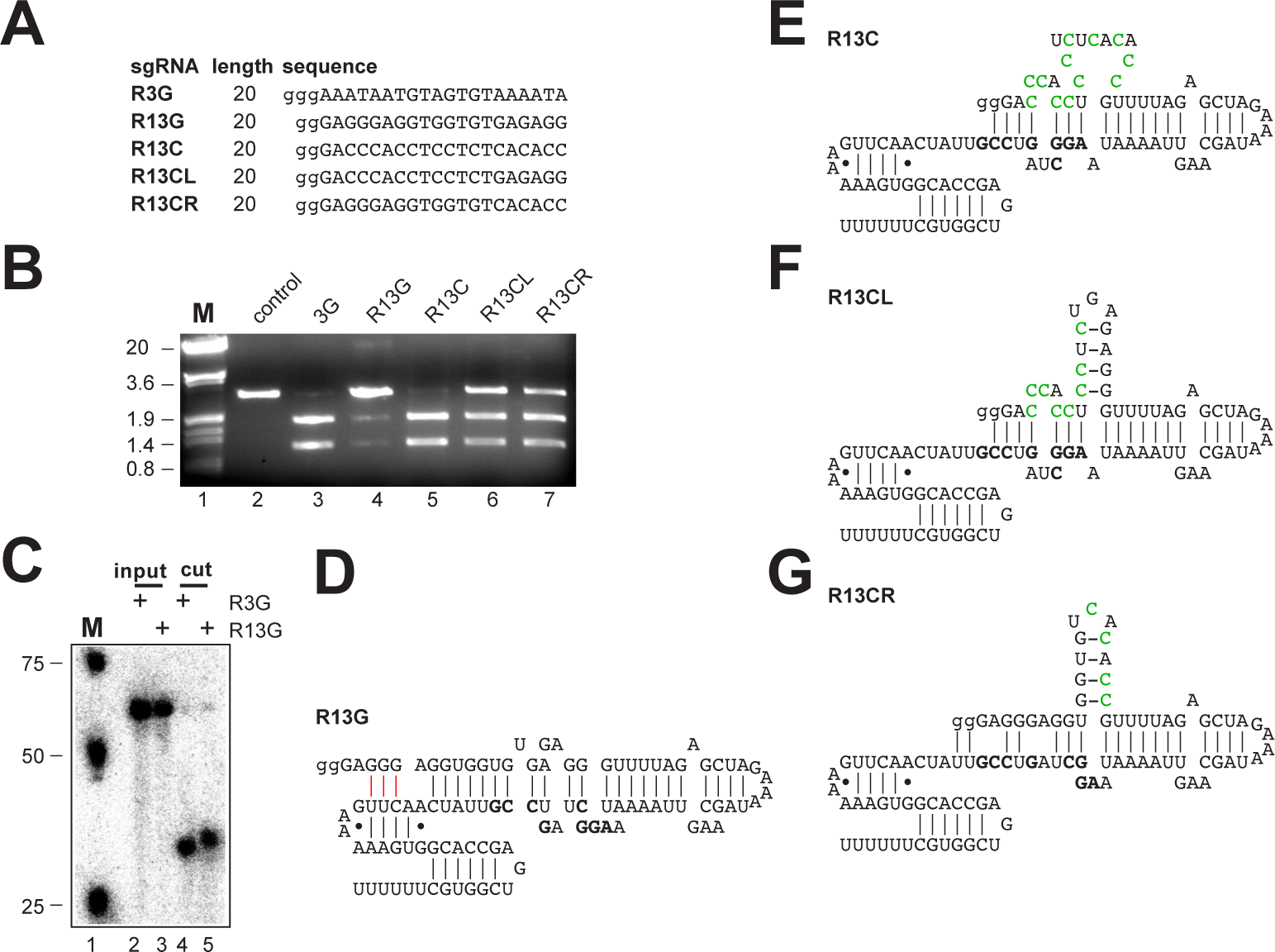
Secondary structure of sgRNAs affects Cas9 mediated cleavage efficiency. (A) Sequences of sgRNAs. Small letter guanosines used for in vitro transcription are not present in the target DNA sequence. (B) Agarose gel showing Cas9 mediated cleavage after 6 h of 3 kb pBS SK+ test-plasmids containing the target sequence with indicated sgRNAs. Plasmids were linearized with ScaI as in the control after Cas9 heat inactivation. Size markers are EcoRI/HinDIII digested λ DNA of 20 kb, 3.6 kb, 1.9 kb and 0.8 kb. (C) Denaturing acrylamide gel showing synthetic oligonucleotides before and after sgRNA/Cas9 mediated cleavage for 1h. (D-G) Structure of sgRNAs. Nucleotides base-pairing in loop 1 are bold. Red lines in D indicate potential base-pairing with nucleotides in loop 2. Green nucleotides indicate mutations compared to sgRNA R13G.

Since the increased number of Gs in sgRNA R13G lead to enhanced base-pairing, we exchanged the Gs with Cs leading to an open structure in the seed region. This sgRNA R13C cleaved the test-plasmid efficiently (**Figs. 2C and 2E**). Introducing Cs in the left or right half of sgRNA R13G lead to short stem loops and inefficient cleavage of the test-plasmid (**Figs. 2C, 2F and 2G**).

To further test to what extent base-pairing impacts on Cas9 activity, we generated sgRNAs L10ds6G and R10ds6GC, where the proximal or the distal half leads to complementary base-pairing of the gRNA with the constant part, respectively (**Figs. 3A,B**). Although both sgRNAs supported Cas9 cleavage of oligonucleotide substrate, the R10ds6GC sgRNA base-pairing with the proximal part was mostly inactive in cleaving the plasmid indicating an impaired ability of Cas9 to scan DNA (**Figs. 3C,D**).

**Figure 3:**
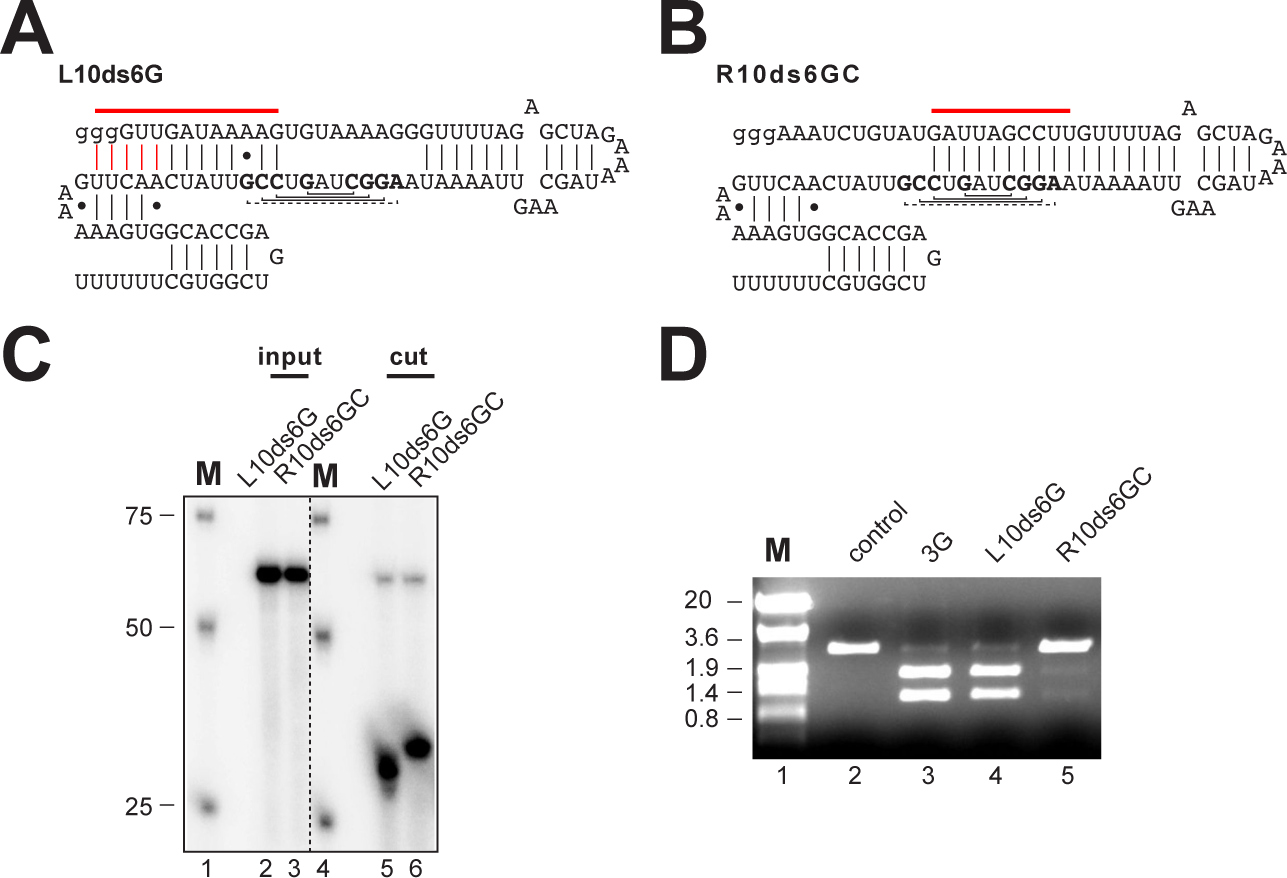
Base-pairing of sgRNAs in the seed-region blocks Cas9 mediated cleavage of a test plasmid. (A, B) Structure of sgRNAs. Horizontal red lines in A and B indicate artificially introduced base-pairing with the sgRNA scaffold, and vertical red lines in a indicate potential base-pairing with nucleotides in loop 2. Nucleotides base-pairing in loop 1 are bold. (C) Denaturing acrylamide gel showing synthetic oligonucleotides before and after sgRNA/Cas9 mediated cleavage for 1h. Note that Cas9 cleavage is heterogeneous. (D) Agarose gel showing Cas9 mediated cleavage after 6 h of 3 kb pBS SK+ test-plasmids containing the target sequence with indicated sgRNAs. Plasmids were linearized with ScaI as in the control after Cas9 heat inactivation. Size markers are EcoRI/HinDIII digested λ DNA of 20 kb, 3.6 kb, 1.9 kb and 0.8 kb.

Taken together, these results demonstrate that the structure of the sgRNA is important for efficient Cas9 mediated DNA scission in vitro. Furthermore, high G content and base-pairing in the distal part of the gRNA also impairs DNA scission, while base-pairing in the proximal part is tolerated. To further substantiate these findings, we analyzed the structures of sgRNAs from previous studies in mammalian cells and *Drosophila* with regard to their cleavage efficiency of previous attempts to define rules for sgRNA cleavage efficiency in vivo(Ren et al. 2014; Graf et al. 2019). Indeed, in 39 sgRNAs designed for use in *Drosophila* reduced cleavage efficiency in nine sgRNAs is associated with disturbances of the sgRNA secondary structure resulting in a cleavage efficiency below 35% (**Supplemental Figs. S3 and S4, Supplemental Table S1C**)(Ren et al. 2014). Similarly, from 22 sgRNAs designed for use in mammalian cells, 13 had a cleavage efficiency below 35% associated with disturbances of the sgRNA secondary structure (**Supplemental Fig S5, Supplemental Table S1D**)(Graf et al. 2019). Similar result were also observed for the efficiency of sgRNAs in honey bees(Roth et al. 2019).

Given the requirement for correct folding of the sgRNA for efficient Cas9 mediated DNA scission, we further examined the X-ray crystal structure of the Cas9-sgRNA-DNA complex to see whether this would provide additional instructions to design sgRNAs (Nishimasu et al. 2014). Indeed, the first two nucleotides, adenosine 51 and 52 (A_51_ and A_52_, **Supplemental Figs. S6 and S7**) after the tetraloop form an aromatic base-stacking interaction with phenylalanine 1105 (Phe_1105_) of Cas9. Furthermore, these interactions are stabilized by guanosine 62 (G_62_) forming non-Watson Crick hydrogen bonds with A_51_ A_52_ and Phe_1105_, and uracil 63 (U_63_) forms a base-stacking interaction with A_52_. These interactions indicate that base-pairing of the gRNA with these nucleotides of the constant part of the sgRNA reduce Cas9 activity in Cas9 cleavage assays in vitro and mutagenesis in vivo.

### Large scale evaluation of novel sgRNA design parameters

Next, we incorporated all the features previously published and from this study into a bioinformatic sgRNA design tool, https://platinum-crispr.bham.ac.uk/predict.pl (Supplemental Code)(Hsu et al. 2013; Doench et al. 2014; Gagnon et al. 2014; Ren et al. 2014; Wang et al. 2014; Chari et al. 2015; Farboud and Meyer 2015; Moreno-Mateos et al. 2015; Doench et al. 2016; Liu et al. 2016; Abadi et al. 2017; Chuai et al. 2018; Labuhn et al. 2018; Graf et al. 2019; Zhang et al. 2019; Michlits et al. 2020; Sledzinski et al. 2020). The included features include validation of intact secondary structures (tetraloop, loop 2 and 3, **Figs. 1-3**), presence of a tetraloop bulge mimic (**Fig. 1K**), self-complementarity of the gRNA (nts 1-20, **Figs. 2F,G**)(Moreno-Mateos et al. 2015; Thyme et al. 2016; Jensen et al. 2017), GC content in the six nucleotide seed region of the gRNA (nts 15-20, **Supplemental Table S1A, S1C and S1D**, (Ren et al. 2014; Wang et al. 2014; Graf et al. 2019), GC content of the gRNA (nts 1-20, **Supplemental Table S1A, S1C and S1D**, (Ren et al. 2014; Wang et al. 2014; Graf et al. 2019), the UUYY motif (nts 16-20), which results in complete base-pairing (**Fig. 3**) and can act as a Pol III termination signal(Gao et al. 2018), the UCYG and CYGR motifs (nts 16-20) associated with lower cleavage efficiency (Graf et al. 2019) and lack of base-pairing of nucleotides 40, 41, 51 and 52 that are engaged in contacts with Cas9 (**Supplemental Figs. S6 and S7**).

PlatinumCRISPr operates at a cut-off prediction for either low or high efficiency, which is based on whether sgRNA folding is correct or whether certain detrimental motifs are present, but also on sequence composition. Although a score could be implemented for sequence composition, in practice, cleavage efficiency increases within an narrow range making a score at this point not feasible.

The sequence of any organism can be used with the PlatinumCRISPr server as long as the genome is sequenced (e.g. accessible via the Ensembl Genome browser). For the most common model organisms, the PlatinumCRISPr server allows to choose the organism and will calculate off targets (additional hits in the genome) as complete match for the 16 nucleotides before the tracrRNA as this is the minimal sequence for DNA scission (Fig. 1G). For other organisms off target effects need to be determined manually by BLAST, e.g. using the NCBI webserver.

Although DNA scission is completely abrogated by single mismatch to the target gene in *Drosophila* (Ren et al. 2014), off target scission has been reported in tissue culture cells (Fu et al. 2013; Hsu et al. 2013). To compare off target effects with sgRNA cleavage efficiencies predicted by either PlatinumCRISPr, DeepSpCas9 or Azimuth (standard scoring in CHOPCHOP) we used a large scale double-strand break analysis from Cas9 expression across 59 targets using tagmentation-based tag integration site sequencing (TTISS)(Schmid-Burgk et al. 2020). In this analysis comparing off target effects predicted by Platinum CRISPr or CHOPCHOP, we find that off target effects measured in tissue culture cells increase with predicted cleavage efficiency for PlatinumCRISPr (one-sided Wilcoxon Rank-Sum test p=0.009) and DeepSpCas9 (Spearman’s rho, p=0.0016), but not for CHOPCHOP (rho 0.1, p=0.41).

To identify additional parameters affecting sgRNA cleavage efficiency, we performed a motif analysis among the 35% low scoring sgRNAs for a number of different data sets (Doench et al. 2014; Gagnon et al. 2014; Ren et al. 2014; Wang et al. 2014; Chari et al. 2015; Farboud and Meyer 2015; Hart et al. 2015; Moreno-Mateos et al. 2015; Varshney et al. 2015; Xu et al. 2015; Doench et al. 2016; Gandhi et al. 2017; Xiang et al. 2021), but we did not find motifs associated with low performance in individual data sets. In particular, we found no evidence for 4 T’s to be enriched that could act as a termination signal for RNA Pol III (Gao et al. 2018; Graf et al. 2019). We then analysed the performance of a *Drosophila* sgRNA data set (Ren et al. 2014) according to sgRNA design parameters described above. Our design tool PlatinumCRISPr selected 12 from 39 sgRNAs and those showed a cleavage efficiency of 58 % or more (**Fig. 4A**). We then analysed a number of sgRNA prediction tools for this data set including Chariscore (Chari et al. 2015), Crispron (Xiang et al. 2021), DeepSpCas9 (Kim et al. 2019), DoenchScore (Doench et al. 2014), Azimuth (implemented in ChopChop)(Doench et al. 2016), Moreno-Mateos Score (Moreno-Mateos et al. 2015), Wang Score (Wang et al. 2014), Wong Score (Wong et al. 2015) and Xu Score (Xu et al. 2015). PlatinumCRISPr significantly outperformed all of these prediction tools with the *Drosophila* data set (Ren et al. 2014)(**Fig. 4B**).

**Figure 4:**
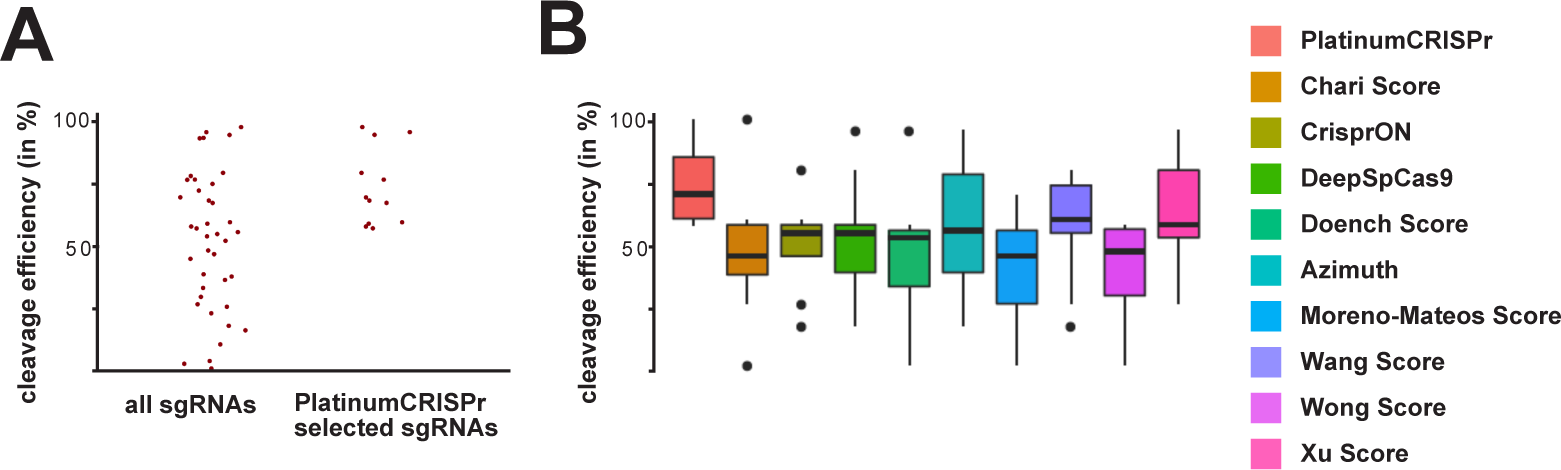
PlatinumCRISPr selects high efficiency sgRNAs and outperforms other sgRNA selection tools for a *Drosophila* data set. (A) Comparison of the in vivo efficiency of all sgRNAs from the Ren et al. (2014) data set with PlatinumCRISPr selected sgRNAs. B) Comparison of the sgRNA selection performance of PlatinumCRISPr with other sgRNA selection tools.

Next, we analysed 14 data sets from various organisms (*Drosophila*, zebrafish, sea squirt, worms and cell culture cells,), which determined sgRNA cleavage efficiency for their performance using the PlatinumCRISPr design tool (Doench et al. 2014; Gagnon et al. 2014; Ren et al. 2014; Wang et al. 2014; Chari et al. 2015; Farboud and Meyer 2015; Hart et al. 2015; Moreno-Mateos et al. 2015; Varshney et al. 2015; Xu et al. 2015; Doench et al. 2016; Gandhi et al. 2017; Xiang et al. 2021). For overall performance (**Fig. 5**), six data sets yielded significant (p≤0.05) enrichment of high efficiency performing sgRNAs (Ren et al. 2014; Wang et al. 2014; Chari et al. 2015; Moreno-Mateos et al. 2015; Xiang et al. 2021), and five showed enrichment (p≤0.25)(Gagnon et al. 2014; Farboud and Meyer 2015; Hart et al. 2015; Varshney et al. 2015; Xu et al. 2015), while two failed to show enrichment for most of the parameters (Doench et al. 2014; Doench et al. 2016). In this analysis, we noticed that structural constraints were significantly more important in cold-blooded organism, where sgRNAs delivery is by injection in the absence of selection in contrast to cell culture cells, where delivery is by transfection and selection for chronic exposure to sgRNAs for up to 10 days before analysis.

**Figure 5:**
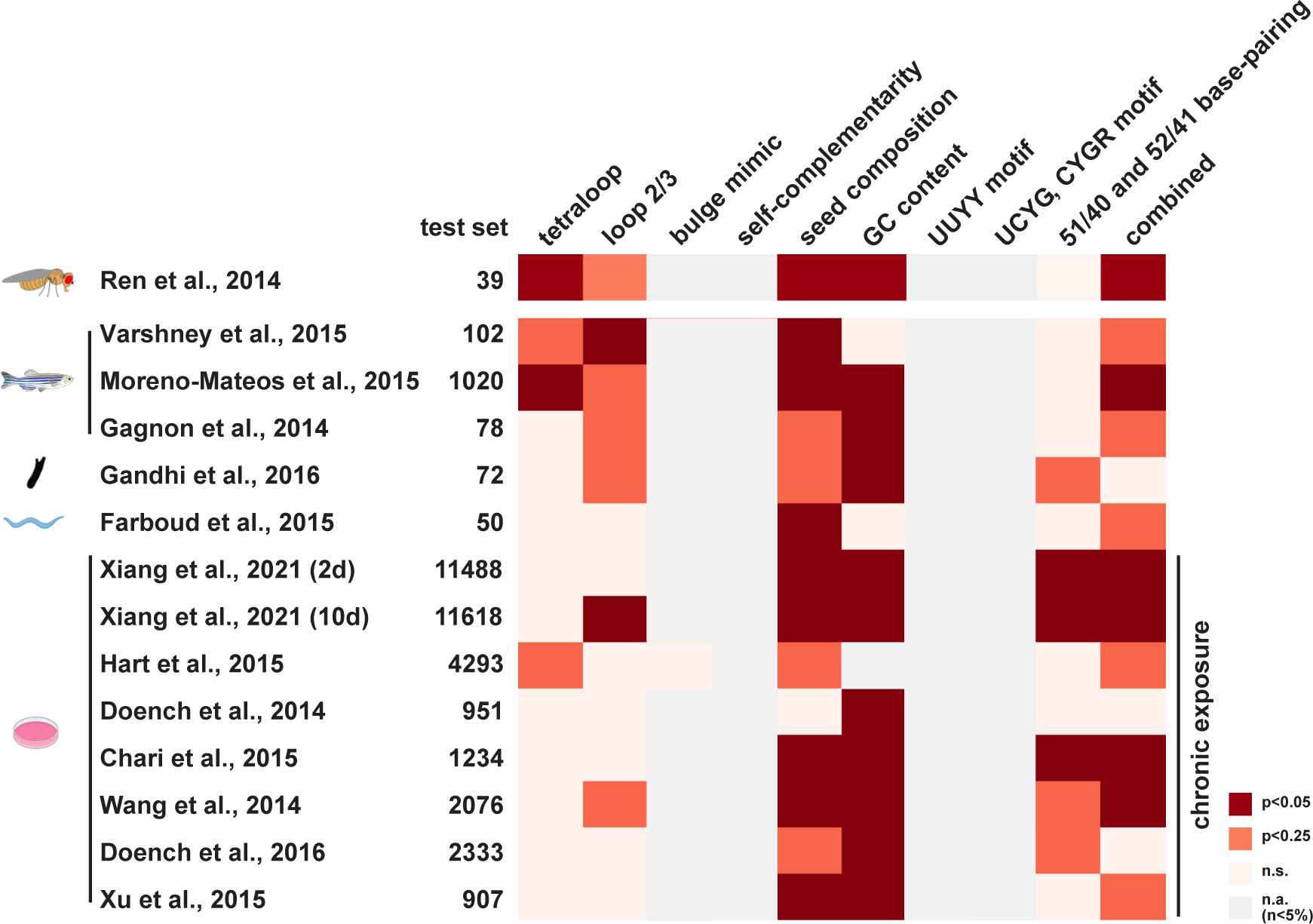
Comparison of individual sgRNA selection criteria for performance with different data sets. The different data sets with the number of sgRNAs tested are indicated on the left. The different selection criteria are shown on top. Significant and enriched performances are indicated in red and orange, respectively (p<0.05 and p<0.25). Criteria with numbers below 5% are indicated in beige, and criteria already applied to the data set are shown in grey.

When we analysed the performance of PlatinumCRISPr, Chariscore (Chari et al. 2015), Crispron (Xiang et al. 2021), DeepSpCas9(Kim et al. 2019), Doench Score (Doench et al. 2014), Azimuth (implemented in ChopChop)(Doench et al. 2016), Moreno-Mateos Score (Moreno-Mateos et al. 2015), Wang Score (Wang et al. 2014), Wong Score (Wong et al. 2015), Xu Score (Xu et al. 2015) and CRISPick (Sanson et al. 2018) prediction tools (**Supplemental Figs. 8A-L**) with the different data sets determining sgRNA cleavage efficiency we found that Wang Score performed best on the Doench data set and that PlatinumCRISPr and Moreno-Mateos performed best with *Drosophila* and zebrafish generated data sets, respectively, but we found no single prediction tool that stood out. In addition, in five out of six cell culture generated data sets none of the prediction tools outperformed the others or substantially increased prediction efficiency (**Supplemental Figs. 8A-L**).

In addition, we have also analyzed datasets for performance of sgRNA libraries in different cell lines (HeLa, HCT 116 and GBM) (Hart et al. 2015) and find very high correlations for all comparisons (Spearman’s correlation between 0.71 and 0.77) indicative of minimal differences of the cleavage efficiency of sgRNAs in these cell lines. In addition, sgRNAs selected by PlatinumCRISPr or CHOPCHOP show a significantly higher cleavage efficiency than discarded sgRNAs in all three cell lines (p≤0.03).

The DepMap project provides efficacy scores for 193476 gRNAs (between 0 and 1, the median efficacy in the complete data-set is 0.974) (DepMap 2024), of which ∼25% are predicted as cutting by PlatinumCRISPr. A comparison of DepMap gRNAs resulted in a minimally better performance (0.006 %, median efficacy: 0.975) of guides predicted as non-cutting by PlatinumCRISPr, but the biological significance of this observation remains unclear since the overall performance substantially exceeded gRNA performance of other cell-line based experiments with chronic exposure (Supplemental Fig. S8), possibly because gRNAs were preselected (DepMap 2024). As part of this analysis, we noticed that the average cleavage efficiency in the analyzed data sets varied substantially (from 20-75% average cleavage efficiency, **Supplemental Figs. 8A-L**). pointing towards a bias in outcome when testing various prediction tools with different data sets. To identify common patterns among the different prediction tools applied to different data sets in the following analysis we therefor excluded data sets with cleavage efficiencies below 30% for further analysis (Gagnon et al. 2014; Chari et al. 2015; Farboud and Meyer 2015; Doench et al. 2016). The default element of ChopChop is Azimuth (Labun et al. 2019), but also implements elements from Doench Score, Chari Score, Xu Score and Moreno-Mateos Score. We therefore reasoned that a combination of two prediction tools could result in more reliable selection of high efficiency cleaving sgRNAs. When we combined two prediction tools the highest scoring combination was PlatinumCRISPr together with Wong score with an average cleavage prediction of 61% (**Fig. 6A**) and this combination also outperformed in all the remaining data sets (**Fig. 6B**), but this increased the stringency and only very few sgRNAs were selected.

**Figure 6:**
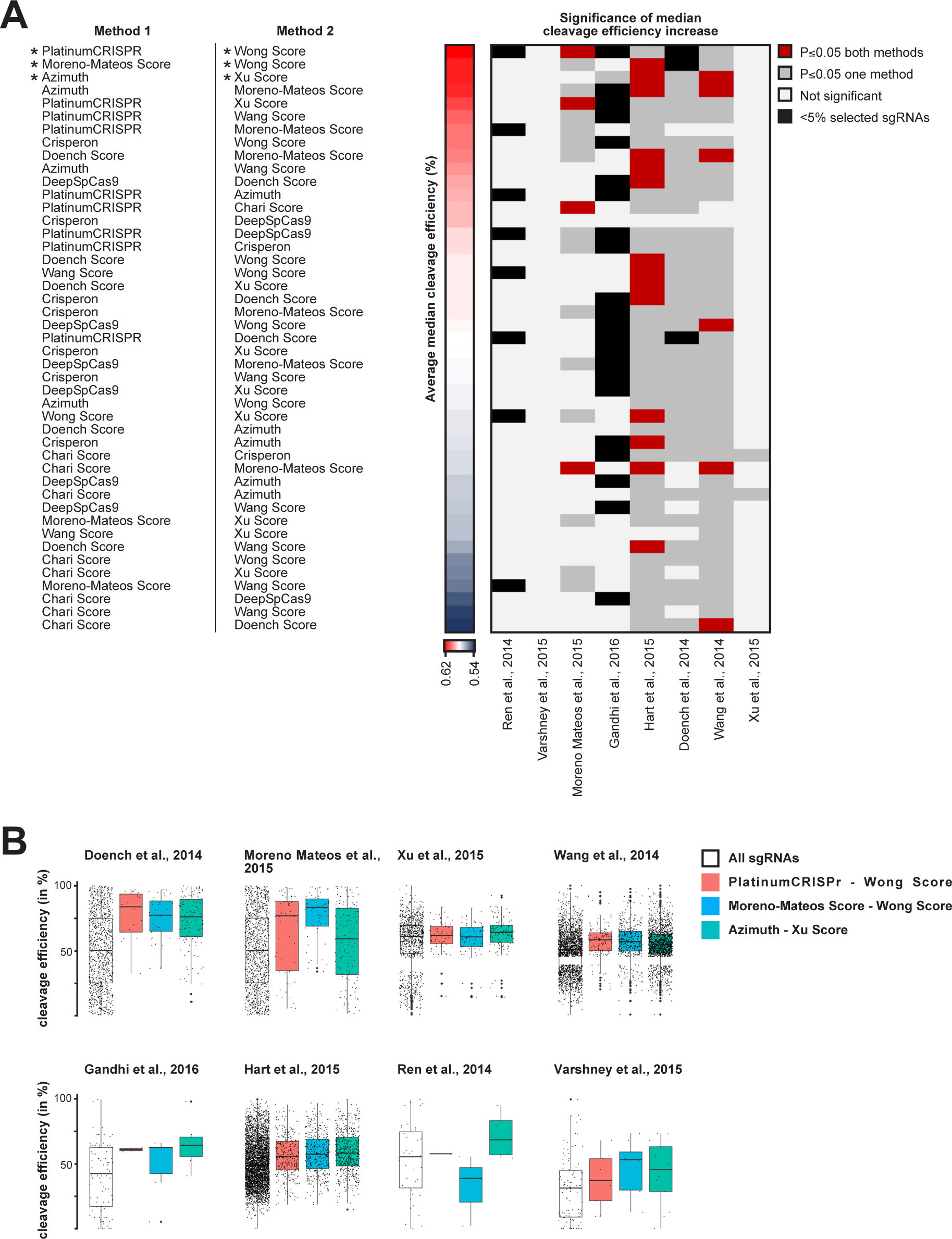
Combinations of two sgRNA selection tools select high efficiency cleaving sgRNAs for several sgRNA efficiency screen data sets. Combinations of sgRNA selection tools in A are listed according to overall cleavage efficiency of selected sgRNA significant (p<0.05) for both methods (red) or one method (dark grey). Black indicates events with less than 5% of sgRNAs selected by at least one method. Comparison of sgRNA selection by different sgRNA selection tools shown as median of the cleavage efficiency for individual data sets (B). The distribution of cleavage efficiencies for all sgRNAs is shown on the left (white box) and for PlatinumCRISPr-Wong Score in red, Wond Score-Xu Score in purple and PlatinumCRISPr-Moreno-Mateos Score in blue.

### A *Drosophila* transformation vector for expression of two sgRNAs

Next, we applied these novel sgRNA design rules to generate *Drosophila* gene deletions. For this purpose we generated a new fly transformation vector with a GFP marker (Solomon et al. 2018), that is easier to select than previously generated *vermillion* marked vectors that require a *vermillion* mutant background for transgene identification(Port et al. 2014; Trivedi et al. 2020). This vector expresses two 20 nt sgRNAs from *U6.1* and *U6.3* promoters (**Supplemental Figs. S9A-C**), harboring a G as first nucleotide as a requirement for expression from the *U6* promoter (Paule and White 2000; Ren et al. 2013). This plasmid can be generated by incorporating the two sgRNA sequences in PCR primers for single step cloning into the plasmid, while previously published vectors require two cloning steps or plasmid recombination (Port et al. 2014; Trivedi et al. 2020). This sgRNA vector can then be injected into *Drosophila* expressing Cas9 in the germline for CRISPR to induce mutations. Alternatively, this vector can be used to generate a transgenic line via the attB site using phiC31 integrase mediated transformation. This fly strain is then crossed to a line expressing Cas9 in the germline for generation of the desired genetic lesion. Transgenically provided sgRNA/Cas9 generally results in a higher efficiency, because the sgRNAs are been provided maternally. Moreover, excision efficiency can be further increased by mutagenesis over a chromosomal deficiency which will removing the repair template for homologous recombination.

### Efficient generation of gene deletions by sgRNA/Cas9 using transposon markers

We then applied the PlatinumCRISPr tool to identify high efficiency sgRNAs to generate gene deletions in *Drosophila Ythdc1* and *Ythdf*, that bind to *N*^6^ methylated adenosines (m^6^A) in mRNA important in development (Dezi et al. 2016; Haussmann et al. 2016; Roignant and Soller 2017; Balacco and Soller 2019; Anreiter et al. 2021). *Ythdc1* and *Ythdf* are located on the third chromosome and transposon inserts *Mi{MIC}YT521-B^MI02006^* and *PBac{SAstopDsRed}^LL04081^* marked with *GFP* or *RFP*, respectively, are available from stock centers. They were combined with an X-linked *vasCas9* or *nosCas9* for germline expression of Cas9. To allow for detection of transposon loss in the YTH protein genes the GFP and RFP markers of the *vasCas9* insert had been removed. These flies were then crossed to the GFP marked sgRNA construct inserted on the 3^rd^ chromosome (**Figs. 7A and F**). The sgRNA insert has a weak GFP marker and can generally be distinguished from *Mi{MIC}YT521-B^MI02006^*inserts. Females from this cross were mated with males containing *TM3 Sb/TM6 Tb* double-balancers in single crosses to recover the sgRNA induced individual deletions and avoid analysis of clonal events. The male progeny was then screened for loss of the GFP or RFP marker. Males which had lost the marker were detected in 100 % (n=9) of the crosses for *Ythdc1* and in 88 % (n=9) for *Ythdf*, respectively. This is a substantial increase of efficiency over imprecise P-element excision with a frequency of 0.01 to 1% (Soller et al. 2006; Haussmann et al. 2016; Haussmann et al. 2022). In those crosses with loss of markers, all males for *Ythdc1* had lost the GFP marker, while for *Ythdf* the average frequency of marker loss was 42 % (n=8). In the reverse cross for *Ythdf*, where sgRNA/Cas9 mediated excision is induced in males, the frequency was 0% (n=5) indicating that the low frequency in males is linked to the absence of recombination in males. The identified single marker-less males were then crossed to *TM3 Sb/TM6 Tb* double-balancers to establish a line and analysed with PCR using primers next to the deletion breakpoints yielding a short PCR product (**Figs. 7B and G**). A PCR product had been obtained in all lines were the marker had been lost indicating that the expected deletion had indeed been generated. To generate *Ythdc1* excision lines *nosCas9* was used for germline expression of Cas9 as *vasCas9* together with sgRNAs targeting *Ythdc1* resulted in female sterility.

**Figure 7:**
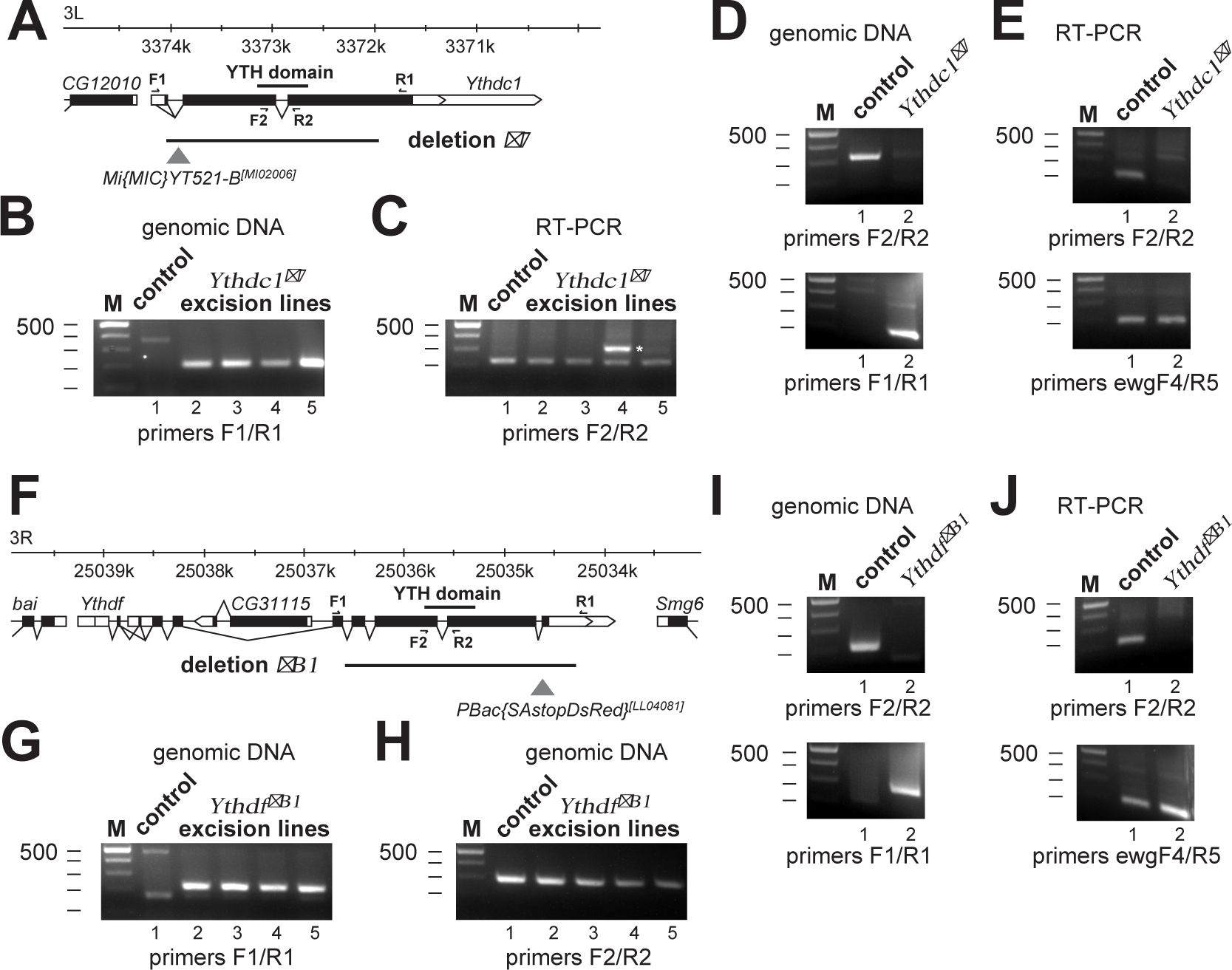
Generation of gene deletions in *Drosophila* YTH protein genes using two sgRNAs/Cas9 and transposon markers. (A) Schematic to the *Ythdc1* locus indicating transcripts (white boxes) and the ORF (black boxes) below the chromosome. Primers used are indicated on top and below the transcripts. The w+ marked transposon used for detecting deletions in the locus is indicated by a triangle and the deletion generated is indicated by a line. (B) Agarose gels showing PCR products amplified from genomic DNA of control or *Ythdc1* transposon excision lines using primers flanking the deletion. Presence of a PCR product indicates the expected gene deletion. DNA markers are indicated on the left. (C) Agarose gels showing RT-PCR products amplified from cDNA of control or *Ythdc1* transposon excision lines using internal primers flanking an intron. Ectopic insertion in the opposite orientation is indicate by an asterisk (lane 4) as in this instance the transcript is not spliced. DNA markers are indicated on the left. (D) Agarose gels showing PCR products amplified from genomic DNA of control or *Ythdc1Δ^7^* flies using internal primers and primers flanking the deletion. DNA markers are indicated on the left. (E) Agarose gels showing RT-PCR products amplified from cDNA of control or *Ythdc1Δ^7^* flies using internal primers in *Ythdf* and *ewg* genes. The PCR product of the *ewg* gene was used as loading control. DNA markers are indicated on the left. (F) Schematic to the *Ythdf* locus indicating transcripts (white boxes) and the ORF (black boxes) below the chromosome. Primers used are indicated on top and below the transcripts. The RFP marked transposon used for detecting deletions in the locus is indicated by a triangle and the deletion generated is indicated by a line. (G) Agarose gels showing PCR products amplified from genomic DNA of control or *Ythdf* transposon excision lines using primers flanking the deletion. Presence of a PCR product indicates the expected gene deletion. DNA markers are indicated on the left. (H) Agarose gels showing PCR products amplified from genomic DNA of control or *Ythdf* transposon excision lines using primers flanking an intron. DNA markers are indicated on the left. (I) Agarose gels showing PCR products amplified from genomic DNA of control or *YthdfΔ^B1^* flies using internal primers and primers flanking the deletion. DNA markers are indicated on the left. (J) Agarose gels showing RT-PCR products amplified from cDNA of control or *YthdfΔ^B1^* flies using internal primers in *Ythdf* and *ewg* genes. The PCR product of the *ewg* gene was used as loading control. DNA markers are indicated on the left.

### sgRNA/Cas9 scissioned fragments insert elsewhere in the genome

After establishing the lines, we noticed that all lines (n=10) established for the *Ythdc1* deletion did not show the flightless phenotype previously reported(Haussmann et al. 2016). Therefore, we selected four lines for further analysis by RT-PCR from RNA (*Ythdc1* excision lines, **Fig. 7C**) or of genomic DNA (*Ythdf* excision lines, **Fig. 7H**) of homozygous flies with primers that were within the deletion and also flanked an intron. Unexpectedly, a copy of the gene was still present in all *Ythdc1* and *Ythdf* excision lines analyzed suggesting that the deleted fragment had been inserted elsewhere in the genome.

To remove this ectopic insert(s), positive lines were crossed to *w+* marked deficiencies and out-crossed for two generations. The X and 2^nd^ chromosomes were then exchanged to establish null-mutant lines from single chromosomes for *Ythdc1* and *Ythdf* that were confirmed by RT-PCR to be free of any ectopic inserts (**Figs. 7D,E and Figs. 7I,J**). To avoid such complications in the future we generated a *PBac w+* containing vector which can be efficiently inserted into a locus by cloning left and right homology arms either to induce a partial deletion upon insertion. Alternatively, mutations in sgRNA cleavage sites can be introduced into the homology arms to insert point mutations. Afterwards, the *PBac w+* is removed scar less by a transposase.

## Discussion

### sgRNA structure and sequence composition contribute to sgRNA/Cas9 DNA scission efficiency

DNA scission by the sgRNA/Cas9 complex is highly specific and requires complete base-pairing between the sgRNA and the target DNA generally not tolerating single miss-matches(Ren et al. 2014; Farboud and Meyer 2015). This feature makes the sgRNA/Cas9 complex an ideal tool for genome editing, but its use is currently limited by the low predictability to cut its target in the genome(Haeussler et al. 2016; Labuhn et al. 2018; Sledzinski et al. 2020).

Here, we discovered that the structure of the sgRNA is a key determinant for the scission efficiency of the sgRNA/Cas9 complex in *Drosophila.* However, only about 50% of sequences adjacent to PAM sites constitute sgRNAs that fold properly or are not compromised by unfavorable base-pairing. In support of these two levels of interference, sgRNA R13GC correctly folds the tetraloop and loop2/3, but did not cleave a short oligonucleotide substrate. This indicates that the structure of this sgRNA blocks the catalytic activity of the sgRNA/Cas9 complex, likely by mimicking the bulge structure of the tetraloop. Second, some sgRNAs allowed cleavage of short oligonucleotide substrates (e.g. L7GC and R10ds6GC), but did not support efficient DNA scission of the target sequence in the context of a 3 kb test-plasmid. Likely, these sgRNAs interfere with the ability of the sgRNA/Cas9 complex to scan the DNA for target sites.

When we analyzed cleavage efficiencies of sgRNAs used in *Drosophila* (Ren et al. 2014), we observed a good overlap with the ability of those sgRNAs to adopt the correct structure and having a high cleavage efficiency. Further refinement to the design of sgRNAs comes from the recognition that the GC content in the seed region is a major determinant to cleavage efficiency in addition to general GC content. Analysis of the X-ray crystal structure of the Cas9-sgRNA-DNA complex also revealed that the two As in the tracrRNA before the tetraloop engage with Cas9 through base stacking and hydrogen bonds (Nishimasu et al. 2014). Base-pairing of these two As with Us at the end of the sgRNA (N_19_ and N_20_) before the start of the tracrRNA impact sgRNA/CAS9 complex function and reduce cleavage efficiency (Graf et al. 2019). Likewise, if sgRNAs are made by RNA Pol III, termination occurs at the boundary of the tracrRNA if two UU precede the GUUUU of the start of the tracrRNA (Arimbasseri and Maraia 2015; Graf et al. 2019). Folding of the sgRNA is not anticipated to impact on transcription as it occurs afterwards. Motif searches in various dataset to determine sgRNA cleavage efficiencies did not reveal any further motifs that impact on cleavage efficiency. If such bias exists, this would likely have been exploited by parasites of prokaryotic hosts.

### Structural constraints of sgRNAs are more pronounced in cold-blooded animals

The bacterial CRISPR-Cas9 system consists of two RNAs (crRNA and trRNA) that assemble with Cas9 to form the active complex. crRNA and trRNA base-pair through sequence complementarity to form the tetra loop in the active Cas9 complex (Garcia-Doval and Jinek 2017; Jiang and Doudna 2017; Hille et al. 2018). In type II systems the tracrRNA is required for crRNA maturation suggesting that base-pairing takes place while being assembled with Cas9. After the crRNA is trimmed, the entire CRISPR-Cas9 complex can scan genomic DNA for DNA scission sites. Alternatively, the crRNA could hybridize first to genomic DNA and recruit tracrRNA and Cas9 to form a complex for DNA scission on site. In this scenario, the crRNA would not be able to interfere with tracrRNA/Cas9 complex activity by forming an aberrant RNA secondary structure, but whether this second scenario could be applied to more efficient genome editing with reduced off-target cleavage needs to be tested. Of note, the effect of structural constraints are stronger in cold-blooded animals likely reflecting that the optimal temperature for *E. coli* is 37°C. In this context, it would be worth exploring how much longer the sgRNA can be to still support Cas9 DNA scission as longer RNAs would more stably hybridize to DNA. In either case, however, understanding sgRNA/CAS9 complex assembly will inform how to prevent off-target DNA scission.

### Evaluation of different sgRNA prediction tools

In this study, we also compared various sgRNA cleavage efficiency prediction tools with 12 datasets that have determined sgRNA cleavage efficiencies in human cells and various model organisms including *Drosophila*, zebrafish, *C. elegans*, honey bees and seasquirt (Doench et al. 2014; Gagnon et al. 2014; Ren et al. 2014; Wang et al. 2014; Chari et al. 2015; Farboud and Meyer 2015; Hart et al. 2015; Moreno-Mateos et al. 2015; Varshney et al. 2015; Wong et al. 2015; Xu et al. 2015; Doench et al. 2016; Gandhi et al. 2017; Kim et al. 2019; Roth et al. 2019; Xiang et al. 2021). Here, the PlatinumCRISPr tool deemed best for *Drosophila*, Moreno-Mateos Score for zebra fish and Wang Score for one human cell culture screen, but all prediction tools failed to convince for the other five cell culture screens. Possibly, chronic exposure over several days in cell culture systems could lead to a bias in determining sgRNA cleavage, compared to short exposure when injected into early stage embryos like in insects and zebra fish.

Taken together, optimizing sequence composition and structural constraints in sgRNA design essentially contributes to high DNA cleavage efficiency. Accordingly, optimized sgRNAs show very little base-pairing with sequences adjacent to the loop 1 region, or are not complementary to the tetraloop structure and/or the loop2/3 structure. In addition, avoiding base-pairing in the 10 nt seed region prior to the PAM site predicts high efficiency of sgRNAs for DNA scission. Furthermore, avoiding two Us before the PAM site prevents interference with the sgRNA/Cas9 structure.

In any case, introducing the target sequence into a plasmid allows to reliably determine the cleavage efficiency of a particular sgRNA in an vitro assay. However, cellular features such as chromatin state can also impact on Cas9 mediated DNA scission (Singh et al. 2015). These limitations of CRISPR-Cas9 genome editing from biological parameters, including the impact of cellular environment by RNA-binding proteins or chromatin structure, remain to be adequately quantified. In this context, whether genes are expressed at the time of sgRNA cleavage likely also plays a role in the observed cleavage efficiency as expression is associated with less compacted DNA. Accordingly, expressed genes seem to be more efficiently mutagenized than developmentally silenced genes (Port et al. 2014; Trivedi et al. 2020).

Initial concerns about CRISPR-Cas9 specificity were about off target cleavage, but changing one nucleotide in the spacer sequence complementary to the target efficiently abrogates cleavage activity (Ren et al. 2014). PlatinumCRISPr takes off-targets into account by screening the genome for the 16 nucleotides distal of the PAM site, which require an exact match for cleavage, and all four possible PAM sites.

### Limitations for generating gene knock-outs

Generating deletions of entire genes, or essential parts of them is the preferred way to generate a null allele. This approach will avoid complications arising from introducing frameshifts at the beginning of the ORF, as translation could reinitiate from later AUG or CUG start codons (Koushika et al. 1999). In addition, in some genes the RNA has functions on its own, as shown for *oscar* RNA that forms a large RNP particle with Oscar and other RNA-binding proteins at the posterior pole of a *Drosophila* oocyte (Hachet and Ephrussi 2004; Haussmann et al. 2011). Thus, deletion of the entire gene region will discover such additional functions harbored in gene transcripts.

When generating deletions of entire genes, we discovered that the deleted DNA fragment was inserted into the genome and transcribed, resulting in the expression of protein that rescued the flightless phenotype in *Ythdc1* deletion allele. Although the mechanism for the generation of these new inserts is not known, it seemed not to have led to complex chromosomal aberrations, because the inserts could be removed by standard recombination and/or exchange of chromosomes. Retro-transposition has been observed in the *elav* gene, which led to the loss of all introns in *Drosophila* (Samson 2008). Likewise, holometabolous insects generally have three *elav* genes, but honey bees have only one *elav* gene. The honey bee *elav* gene, however, carries features of the other two genes present in *Drosophila*. Hence, *elav* in honey bees could have collapsed in an ancestor from three to one gene by some form of recombination to include parts specific to the other two *elav* genes (Ustaoglu et al. 2021). Likewise, the *amyloid-β precursor protein* (*APP*) gene displays copy number variation in the human brain, which are increased by retro-transposition in sporadic forms of Alzheimer’s disease suggesting evolutionary conserved mechanisms for reinsertion of genomic information (Lee et al. 2018). Re-insertion of fragments cut out by the CRISPR-Cas9 system seems not to be specific to *Drosophila* as it has also been observed in human cells (Geng et al. 2022). Since intron loss and gain occurs during evolution, the mechanism underlying insertion of DNA fragments from sgRNA induced deletions might be responsible for these changes. Thus, when generating sgRNA induced gene deletions, it is essential to test for the absence of any transcripts by RT-PCR, but also to validate a deletion at the DNA level by PCR using flanking primers or Southern blots.

For a more reliable way to generate gene knock-outs, CRISPR-Cas9 induced deletion of only a part of a gene seems more feasible and aligns with previous mutagenesis approaches using marked *P* element transposons (Supplemental information and discussion, **Supplemental Fig. 10A**)(Soller et al. 2006; Haussmann et al. 2016; Bawankar et al. 2021; Haussmann et al. 2022). Such targeted deletions to 5’part of the gene can remove the core promoter consisting of the TATA box and extend into a functional domain (e.g. catalytic domain, DNA-binding domain, RNA recognition motif, etc, **Supplemental Fig. 10B**).

Since human genes are much larger and often have alternative transcription start sites (Soller 2006), deletion of an exon such that splicing of the remaining exons generate a frameshift is a valuable option (**Supplemental Fig. 10C**). Alternatively, a GFP cassette with a poly(A) site can be inserted to terminate the ORF in the beginning, but it needs to be evaluated whether the poly(A) site in the beginning of the gene is used or whether the exon containing the GFP cassette is skipped (Soller 2006; Wierson et al. 2020).

For a more reliable way to generate gene knock-outs in *Drosophila*, we have now developed a *PBac w+* marker that can be inserted when generating a deletion (Dix 2022). In any case, however, a marked locus will allow for rigorous cleaning of the genetic background.

In essence, we have established the rules for designing highly efficient sgRNAs and established methodology to efficiently generate gene deletions. These findings have implications for other RNA based methodologies including prime editing (Anzalone et al. 2019; Bosch et al. 2021).

## Methods

### sgRNA/Cas9 directed DNA cleavage

DNA templates for in vitro transcription were reconstituted from synthetic oligonucleotides. As only the T7 promoter needs to be double-stranded for in vitro transcription, a T7 promoter oligonucleotide (CCTGGC<u>TAATACGACTCACTATA</u>G) was annealed to an anti-sense Ultramer (IDT DNA) encoding the entire sgRNA in addition to the T7 promoter. Alternatively, a 60 nt T7 promoter oligonucleotide with a partial sgRNA was annealed to an anti-sense oligonucleotide encoding the tracrRNA (AAAAAAAGCACCGACTCGGTGCCACTTTTTCAAGTTGATAACGGACTAGCCTTATTTT AACTTGCTATTTCTAGCTCTAAAAC) for 15 min at 40° C (2 µM) and made double-stranded by extension with Klenow fragment of DNA polymerase I according to the manufacturer’s instructions (NEB). Klenow was then heat-inactivated 10 min at 85° C and oligonucleotides were desalted with a G-50 Autoseq Sephadex spin column (GE) before using for in vitro transcription.

Then, sgRNAs were generated by in vitro transcription with T7 polymerase (T7 MEGAscript, Ambion) from synthetic oligonucleotides (0.2 µM) and trace-labeled with ^32^P alpha-ATP (800 Ci/mmol, 12.5 µM, Perkin Elmer) in a 20 µl reaction according to the manufacturer’s instructions.

After DNase I digestion, free nucleotides were removed with a G-50 Probequant Sephadex spin column (GE). Then, sgRNAs were heated for 2 min to 95° C and left at room temperature to adopt folding. Then, sgRNAs were quantified by scintillation counting and analysed on 8-20 % denaturing polyacrylamide gels as described (Dix et al. 2022).

For synthetic substrate DNAs the sense oligonucleotide (1 µM, sgRNA flanking sequences are: TCGAGCATTATATGAAC-sgRNA-GGGTATTGGGGAATTCATTATGC) was labeled with ^32^P gamma-ATP (6000 Ci/mmol, 25 µM, Perkin Elmer) with PNK (NEB). After heat-inactivation of PNK for 2 min at 95° C, sense and anti-sense (anti-sense sgRNA flanking sequences are: GGCCGCATAATGAATTCCCCAATACCC-as sgRNA-GTTCATATAATGC) oligonucleotides were annealed by letting cool down to room temperature and used in sgRNA-Cas9 cleavage assays. For plasmid sgRNA-Cas9 cleavage assays, these annealed oligonucleotides were cloned into a modified *pBS SK+* using a Xho I and Not I cut vector to assay sgRNA/Cas9 activity.

For sgRNA/Cas9 cleavage assays, DNA/sgRNA/Cas9 ratios of 1/10/10 were used in a 10 µl reaction using the buffer supplied (NEB) and DEPC-treated water (Haussmann et al. 2019). Typically Cas9 (100 nM final) was incubated with sgRNA (100 nM) for 10 min at 25° C before adding oligonucleotides (10 nM final) or plasmid DNA (10 nM, corresponds to ∼25 ng/µl final concentration of a 3 kb plasmid). Plasmids were linearized after Cas9 digestion by first heat inactivating Cas9 for 2 min at 95°C, and then adding 10 µl of a restriction enzyme (5 U) in NEB buffer 3. Adding a restriction enzyme together with Cas9 inhibited DNA scission by Cas9. Cleavage of oligonucleotides was analysed on 8 % denaturing polyacrylamide gels and plasmid DNA was analysed on ethidium bromide stained agarose gels.

RNA secondary structure was analyzed with RNAfold at http://rna.tbi.univie.ac.at (Gruber et al. 2008) using the following tracrRNA sequence: GUUUUAGAGCUAGAAAUAGCAAGUUAAAAUAAGGCUAGUCCGUUAUCAACUUGAAAAAGUGGCACCGAGUCGGUGCUUUUUU.

### Criteria for optimal sgRNA design and Cas9 structural analysis

Criteria to select sgRNAs that maintain the structure required for efficient Cas9 DNA scission were implemented in the server accessible at https://platinum-crispr.bham.ac.uk/predict.pl and are as follows. The first nucleotide of the sgRNA needs to be a G for efficient transcription initiation by RNA Pol III (Paule and White 2000). For T7 mediated in vitro transcription, three Gs need to be added to sgRNAs of 23 nucleotides. Disruption of the tetraloop or loop 2 and 3 structures, and the sequence U A/G G C/U A/G of nucleotides 16-20, which will result in a tetraloop bulge mimic, were classified as low efficiency sgRNAs as these parts are recognized by Cas9 (Nishimasu et al. 2014). Similarly, a hairpin loop in the gRNA consisting of 4 or more base pairing nucleotides, or base pairing of nucleotides 17-20 of the gRNA (U/C U/C U U), or base-pairing of eight nucleotides within the seed region with looped-out nucleotides spaced by three base-pairing nucleotides (N11-N20), or a GC content below 15% (1 or 2 nts) or above 50% (11 or more nts) were also considered low efficiency. Medium efficiency was assigned for gRNAs with a GC content of 15-25 % (3-5 nts) or 40-50% (8-10 nts), or a low CG content in the seed region (less than 5 nts in nucleotides 11-20). In addition, two U’s at position 19/20 of the gRNA reduce efficiency because of premature transcription termination (Gao et al. 2018; Graf et al. 2019). Further, we assigned a medium impact if both of the two nucleotides A_51_ A_52_ and G_62_ U_63_ in the three way junction of loop 1 were base paired or seven nucleotides within the seed region (N11-N20) base-paired with looped-out nucleotides spaced by three base-pairing nucleotides. Thus, we deemed an sgRNA optimal to allow Cas9 to cleave DNA with high efficiency, if the GC content is 30-35 % (6-7 nts) and none of the above criteria applied.

Cas9/sgRNA structural complex analysis was done using Chimera as described (Dix et al. 2022).

### RNA extraction, RT-PCR and PCR on genomic DNA

Total RNA was extracted using Tri-reagent (SIGMA) and reverse transcribed with Superscript II (Invitrogen) according to the manufacturer’s instructions using an Oligo(dT) primer. PCR was done for 40 cycles with 1 µl of cDNA, with 1 µg of genomic DNA or from a single fly after freezing and drying in 200 µl of isopropanol. Primers to detect the sgRNA/Cas9 induced deletion in *Ythdc1* were YT F1 (GCCGCTGTGACGCAGAATTTGTGTG) and YT R1 (GGCCGTGCATGTTGCGCATGTAGTCC), and in *Ythdf* were 64F1 (GCCGAGAAAGTGCACAAGGATACGGAG) and 64R1 (CAAGGAATGGCTGAAGCAGACTCCTTG). Primers to amplify parts of the body of the RNA also flanking an intron were for *Ythdc1* YT F2 (CCACGCTGCCGCAGAACGACGCCAATC) and YT R2 (GCGGCAGATCCAGTCAAGCTCGATGAC), and in *Ythdf* were 64F2 (GAGCTGCCTGTCGATTCCCAACTCGTG) and 64R2 (CCGCCCTCTTCGTGTCGCTCCTTGAAG). Primers to amplify parts of the *ewg* gene have been described elsewhere (Koushika et al. 1999).

### Cloning of sgRNAs into *pUC 3GLA U6.1 BbsI*

To clone two sgRNAs expressed by U6 promoters, the “tracrRNA U6.3 promoter” fragment was amplified with left (AAGATATCCGGGTGAACTTC**<u>G</u>**N_19_GTTTTAGAGCTAGAAATAGC) and right (GCTATTTCTAGCTCTAAAA**<u>C</u>**N_19_CGACGTTAAATTGAAAATAGG) sgRNA primers from pUC 3GLA U6.1/3 sgRNA using Pwo polymerase (Roche) with initial 30 sec denaturation at 94°C followed by two cycles 94°C/30 sec, 49°C/40 sec, 72°C/45 sec, then two cycles 94°C/30 sec, 51°C/40 sec, 72°C/45 sec and 22 cycles two cycles 94°C/30 sec, 56°C/40 sec, 72°C/45 sec. Transcription from the U6 promoter initiates with a G (bold, underlined). Although this G does not need to be present in the targeting sequence, it needs be included for folding of the sgRNA. The sequences for the sgRNAs in *Ythdc1* were gACAGGTATTCCCAAACTCAC and GACATGTAGCGTTCCCATGA, and for *Ythdf* were GTCCTGAAATACGAGCACAA and gATAACGAACATGTGGGATCT. The pUC 3GLA U6.1 BbsI vector was cut by BbsI and the “sgRNA1 tracrRNA U6.3 promoter sgRNA2” fragment was cloned by Gibson assembly according to the manufacturer’s instructions (NEB). For sequencing, primer U6.1 Fseq (GCGCGTACGTCCTTCGCATCCTTATG) was used. The sequences for *pUC 3GLA HAi Dscam 3-5, pUC 3GLA U6.1 BbsI* and the *pUC 3GLA U6.1/3 Ythdf sgRNA* have been deposited (MK908409, MK908408 and MK908407).

### *Drosophila* genetics and phiC31 integrase-mediated transgenesis

All *Drosophila melanogaster* strains were reared at 25°C and 40%–60% humidity on standard cornmeal-agar food in 12:12 h light:dark cycle as described (Haussmann et al. 2013). *CantonS* was used as a wild type control. For the *Ythdc1*, the GFP marked *Mi{MIC}YT521-B^MI02006^* transposon insert and the *w+* marked *Df(3L)Exel6094* deficiency were used, and for *Ythdf*, the RFP marked *PBac{SAstopDsRed}^LL04081^* insert and the *w+* marked deficiency *Df(3R)ED6220* were used. For phiC31 mediated transformation, constructs were injected into *y^1^ w* M{vas-int.Dm}ZH-2A; PBac{y+-attP-3B}VK00013* with the landing site inserted at 76A as previously described(Haussmann et al. 2013). Prior to insertion of GFP marked constructs, the GFP and RFP markers had been removed from the *y1 w* M{vas-int.Dm}ZH-2A* landing site by Cre mediated recombination (Bischof et al. 2007; Zaharieva et al. 2015).

### Implementation of PlatinumCRISPr

PlatinumCRISPr is implemented as a Perl script based web-server iteratively evaluating the rule set described in the main text. A guide is classified as “compromised” if any of the rules is violated. For analysis of an sgRNA consisting the target complementary sequence (spacer) and the constant crispr RNA (crRNA) fused to the tracrRNA through an artificial loop is used whereby the first nucleotide of the 20 nt spacer sequence is a G, because a G is needed for transcription (Jinek et al. 2012; Cong et al. 2013). Folding of the sgRNA is computed using RNAFold (Version 2.4.17) and further processed using bpRNA for subsequent interpretation of the dot-bracket code describing the secondary structure by a custom-made Perl script. Notably, the sequence position is calculated from the 3’end of the tracerRNA to allow for variable length of the spacer (between 18 and 23 nt) for custom applications using synthesized RNA consisting of the spacer fused to the crRNA and hybridized to the tracrRNA.

PlatinumCRISPr classifies guides by a binary outcome and typically reports around 70% of sgRNAs as “compromised”. Accordingly, only the top 30% were analysed for the distribution of their reported cleavage efficiency in a given data set for each scoring application (**Supplemental Fig S8).** Statistical significance was calculated using a one-sided Wilcoxon signed-rank test. We are grateful to M. Haeussler for providing published guide sequencing and cleavage efficiency scores (Haeussler et al. 2016).

CrisperON and DeepSpCas9 guide sequencing and cleavage efficiency scores were calculated using published web-interfaces (Wong et al. 2015; Xiang et al. 2021).

For CRISpick analysis, we downloaded pre-calculated on-target score data (options: KO screen, CRISPR mechanism SpyCas9, Hsu TracerRNA and On Target Scoring Rule Set was RS3seq-Chen2013+RS3target) which are available for Mouse, Fly, and Human (Sanson et al. 2018). DepMap inferred guide efficiency data has been downloaded from DepMap (DepMap 2024). For the analysis of sgRNA efficiency in different cell lines (HeLa, HCT 116 and GBM), genes essential in all cells were selected and the calculated cleavage efficiencies were compared by Spearman’s correlation (Hart et al. 2015).

For the analysis of off-target effects, guides from the TTISS data-set with their number of genome wide off-target sites were used (Schmid-Burgk et al. 2020; Chen et al. 2023). For each guide, the number of expected off-targets was computed using Cas-Offinder3 (selecting targets with no bulge, and a minimum of one and a maximum five mismatches) (Hwang et al. 2021). The ratio of measured to expected numbers of off-target sites was then compared between guides predicted to cut by PlatinumCRISPr (Wilcoxon-Rank-sum test) and correlated (Spearman’s correlation) to DeepSpCas9 and CHOPCHOP (Azimuth 2.0 scores) using the crisprScore R-package (Hoberecht et al. 2022).

## Data access

All data used for sgRNA cleavage efficiency analysis to analyze data and the source code for PlatinumCRISPr have been deposited in GitHub (https://github.com/rolandA1234/PlatinumCRISPr) and are also provided as Supplemental Code.tar.gz file.

Sequences for plasmid vectors generated in this study have been submitted to NCBI GenBank (https://www.ncbi.nlm.nih.gov/genbank/about/) with the following accession numbers: MK908409 (*pUC 3GLA HAi Dscam 3-5*), MK908408 (*pUC 3GLA U6.1 BbsI*) and MK908407 (*pUC 3GLA U6.1/3 YTHDF53 sgRNA*).

## Competing interest statement

The authors declare no competing interests

## Acknowledgments

We thank Bloomington and Kyoto stock centers for fly lines, A. Handler for the PiggyBac clone, BacPac for Bac clones and plasmids, the University of Cambridge Department of Genetics Fly Facility for embryo injections for generating transgenic lines, M. Haeusler for datasets and pre-calculated scores, Sung-Eun Yoon from the Korea Drosophila Resource Centre and Ying Tan from Genetivision Corporation for sgRNA efficiency data, Yuan Tian for help with art work and Pawel Grzechnik for comments on the manuscript. For this work we acknowledge funding from the Biotechnology and Biological Science Research Council, the Leverhulme Trust, the MRC DTP to DWJM and a Genetics Society summer studentship to AIH.

## Author contributions

IUH, TCD and MS performed biochemistry and genetic experiments, VD constructed vectors and AH performed genetic experiments. RA performed bioinformatics analysis, programming and server implementation. DWJM analysed data. IUH, RA and MS conceived the project and wrote the manuscript with help from all authors..

## Supplemental Material

### Supplemental information and discussion

#### A guide to generate a gene knockout

Historically, imprecise excision of a marked *P* element transposon represents a versatile approach for directed mutagenesis of a gene in *Drosophila* and other organisms to generate a knock-out as illustrated for a knock-out of *mRNA cap methyltransferase 1* (*CMTr1*) gene (Supplemental Figure 10A) (Soller et al. 2006; Haussmann et al. 2016; Bawankar et al. 2021; Haussmann et al. 2022).

A *P* element transposon (unlike *Minos* or *PiggyBac* elements) can generate deletions by imprecise excision in the presence of a transposase. To increase excision efficiency, this is generally done over a larger chromosomal deficiency (marked with w+, available from stock centres) to prevent repair from the homologous chromosome. This type of widely used mutagenesis has been facilitated by large collections of *P* element insertions in the *Drosophila* genome and availability of corresponding strains from stock centers.

Accordingly, to generate a knock-out of *mRNA cap methyltransferase 1* (*CMTr1*) gene, a *P* element transposon (marked with w+) inserted in the 5’UTR was mobilized and an imprecise excisions were recovered. These deletion were then mapped by inverse PCR and a deletion was selected that removes parts from the 5’UTR extending into the catalytic domain rendering this allele functionally null (Supplemental Figure 10A). Like chemical mutagenesis, imprecise excision of a marked *P* element requires screening for loss of the marker and then for a deletion of the desired extension.

Now, this process is massively facilitated by the use of sgRNA/Cas9 to introduce the desired deletion to generate a gene knock-out. In principal, deletions can be recovered after injection of a plasmid expressing two sgRNA under the *U6* promoter into *Drosophila* embryos expressing Cas9 under a germline promoter. To increase efficiency, however, we advise to use a transposon marker in the gene. In addition, to prevent repair from the homologous chromosome, the transposon is best excised in the presence of a marked larger chromosomal deficiency (available from stock centres). In this configuration, however, only 25-50% of embryos will have the desired genotype, because of lethality of the deficiency chromosome and potentially the transposon insert. Hence, a transgene expressing two sgRNA under the *U6* promoter facilitates the procedure, but will take longer.

To generate a knock-out for a typical *Drosophila* gene by sgRNA/Cas9, parts of the promoter (indicated by the TATA box) and a main functional domain (e.g. a catalytic domain, an RNA Recognition Motif, a zinc-finger DNA binding domain, etc) are deleted. Accordingly, gRNAs are then selected in these parts of the gene by entering the relevant sequence into PlatinumCRISPr (dashed lines, Supplemental Figure 10B).

Potential deletions are first identified by loss of the transposon marker and then validated by PCR using primers flanking the deletion. If a deletion is present, a short PCR product will be amplified, while a PCR for the parental chromosome will be much larger and likely unproductive. To check for re-insertion of the deleted fragment, PCR with primers flanking an intron can be used. If the deleted fragment reinserted in the sense orientation and is expressed, a shorter fragment will be detected by RT-PCR compared to a larger fragment when the insert is expressed from the anti-sense strand.

Human genes are on average ten times larger than *Drosophila* genes due to more and larger introns (Soller 2006). Typically, human genes have short exons of around 150 nucleotides which are separated by large introns. These features require adaptation of the knock-out approach.

One possibility to overcome this challenge of increased gene size is to remove the promotor together with the first exon. However, this is often not feasible because most human genes have several transcription start-sites, physically separated by several kilobases.

An alternative approach is to delete an exon such that splicing will generate a frame-shift. To identify suitable gRNAs to delete this exon, the sequences before and after are entered into PlatinumCRISPr. To avoid splicing complications, it is best to avoid gRNAs in splice sites (select gRNAs before the branchpoint and about 30 nucleotides after the 5’ splice site, Supplemental Figure 10C).

Such a deletion can conveniently be monitored by RT-PCR with primers in the flanking exons as they generate a short PCR product if the desired deletion is present, but a longer PCR for the parent. If sgRNA/Cas9 induced deletions are made in a population of cells (e.g. under puromycin selection), sgRNA/Cas9 efficiency can be monitored this way through the presence of two PCR products (with and without the deleted exon).

**Supplemental Figure S1.**
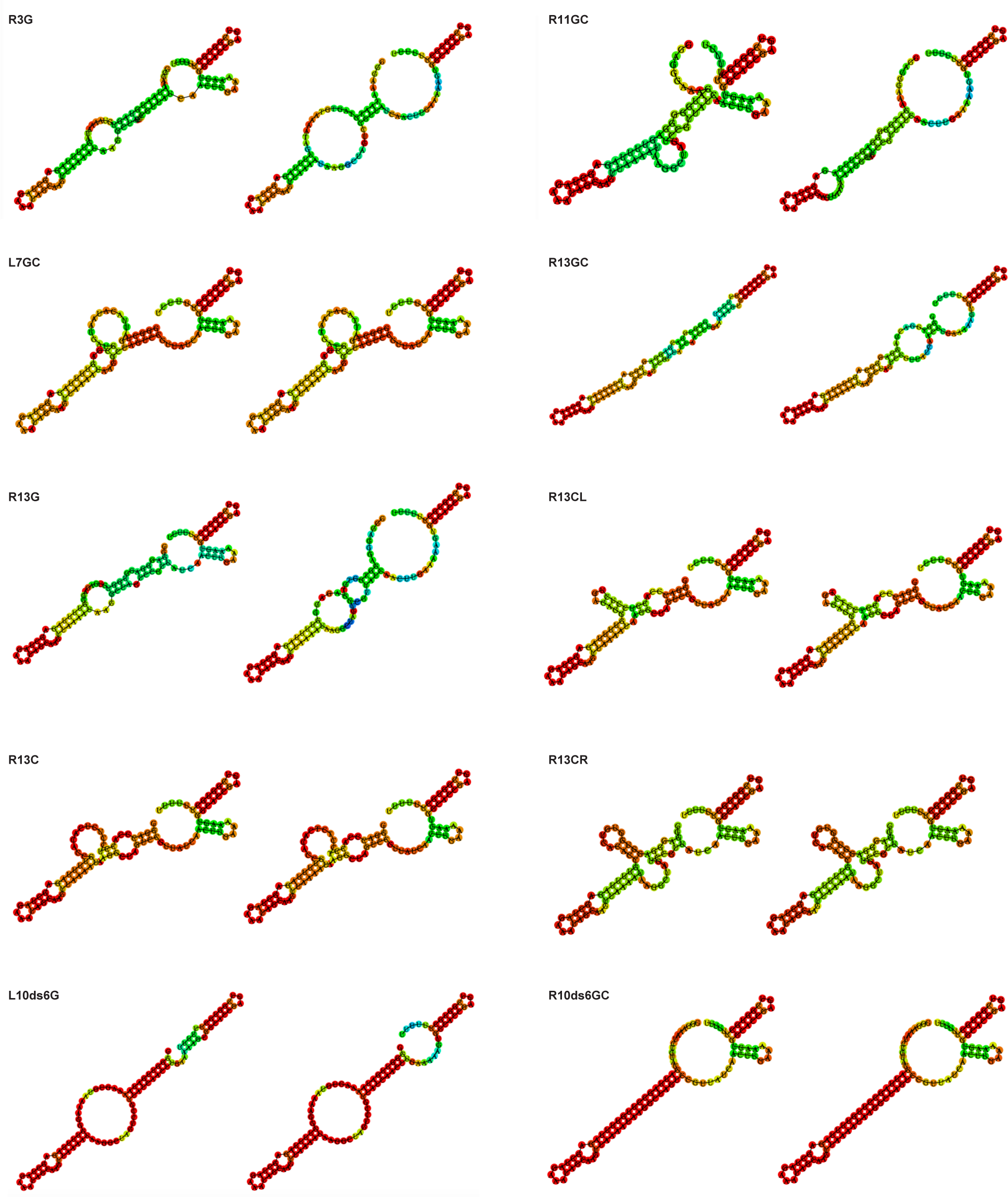
Secondary structures of sgRNAs used in Figs 1-3 predicted by RNAfold.

**Supplemental Figure S2.**
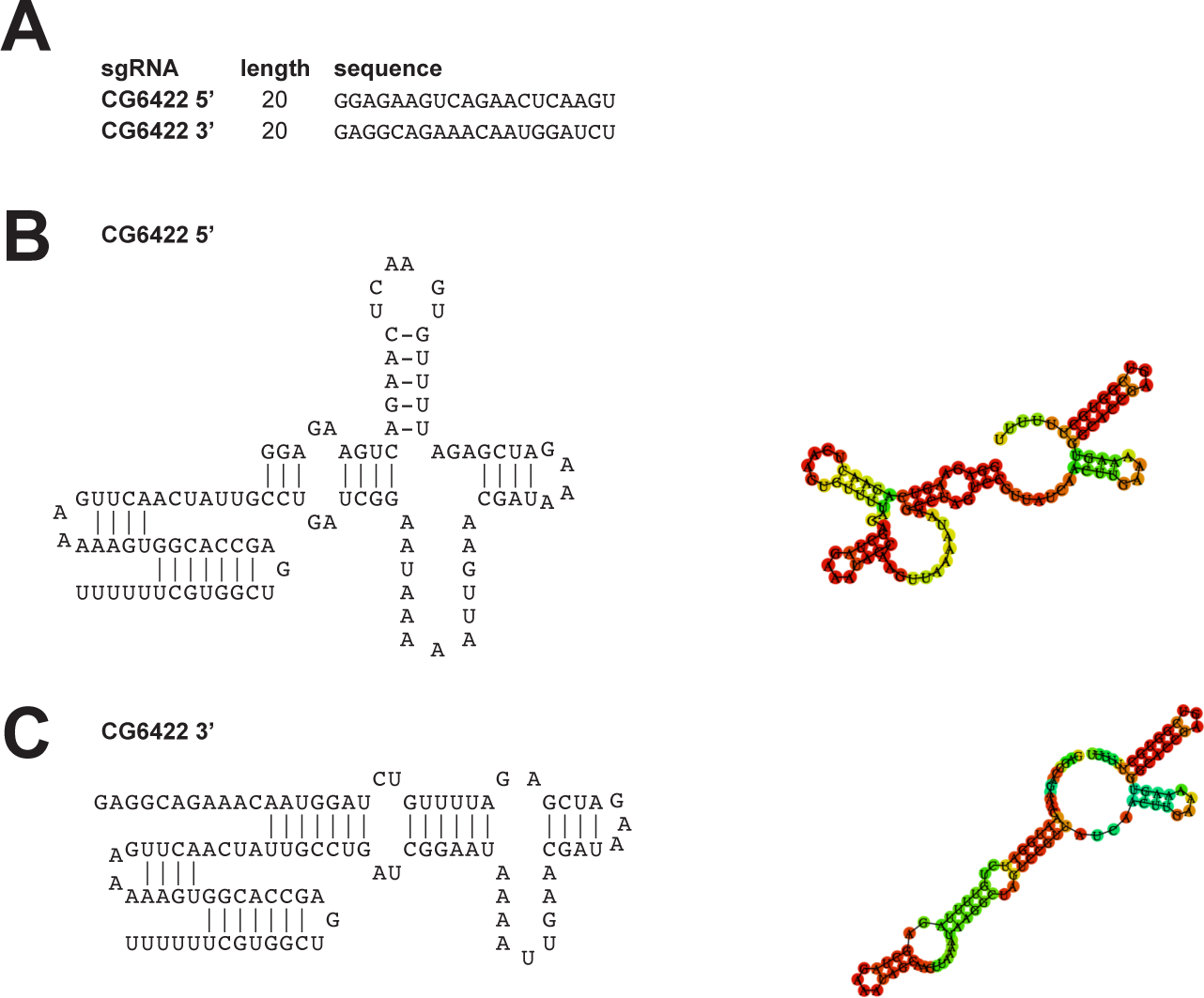
Inactive sgRNAs targeted to the Drosophila ythdf (CG6422) gene. **(A)** Sequences of sgRNAs targeting the CG6422 locus for generating a deletion. **(B, C)** Secondary structures of sgRNAs. The scaffold is shown on the left indicating Watson-Crick base-pairing by lines and the predicted secondary structure by RNAfold is shown on the right.

**Supplemental Figure S3.**
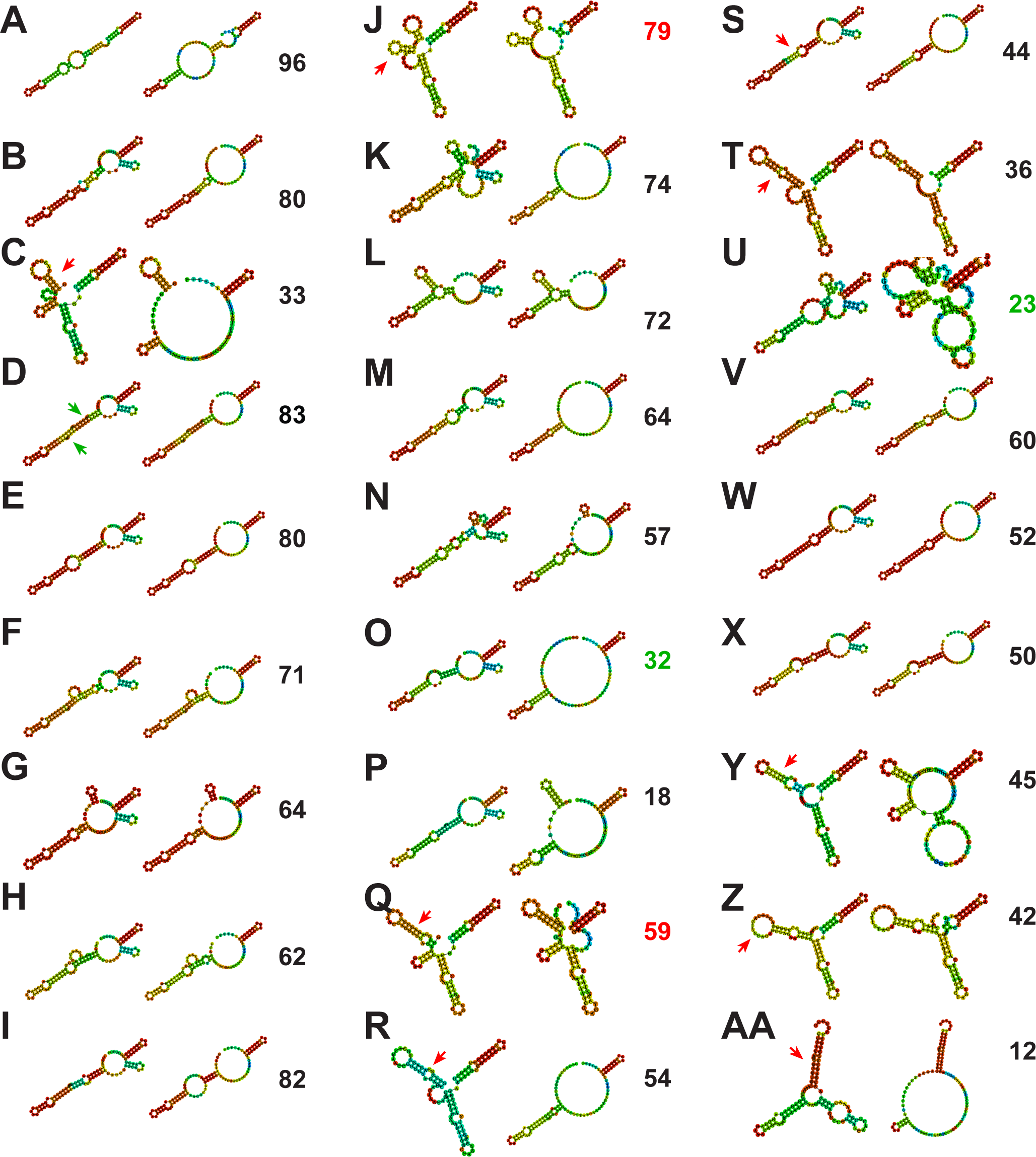
Secondary structure of sgRNAs targeted to the Drosophila white gene from Ren et al. (2014). **(A-AA)** Secondary structures of sgRNAs predicted by RNAfold. Minimal energy and proximity base-pair structures are shown on the left and right, respectively. Red and blue indicated high and low probabilities for the adopted structural base-pairing assignment, respectively. The number on the right indicates the effectiveness of inducing heritable mutations after injection into fly embryos as determined by Ren et al. (2014). Red and green numbers indicate an effectiveness which is too high or too low compared to predicted DNA cleavage efficiency. Red arrows point towards structural features likely limiting DNA cleavage and green arrows point towards structural features supporting DNA cleavage.

**Supplemental Figure S4.**
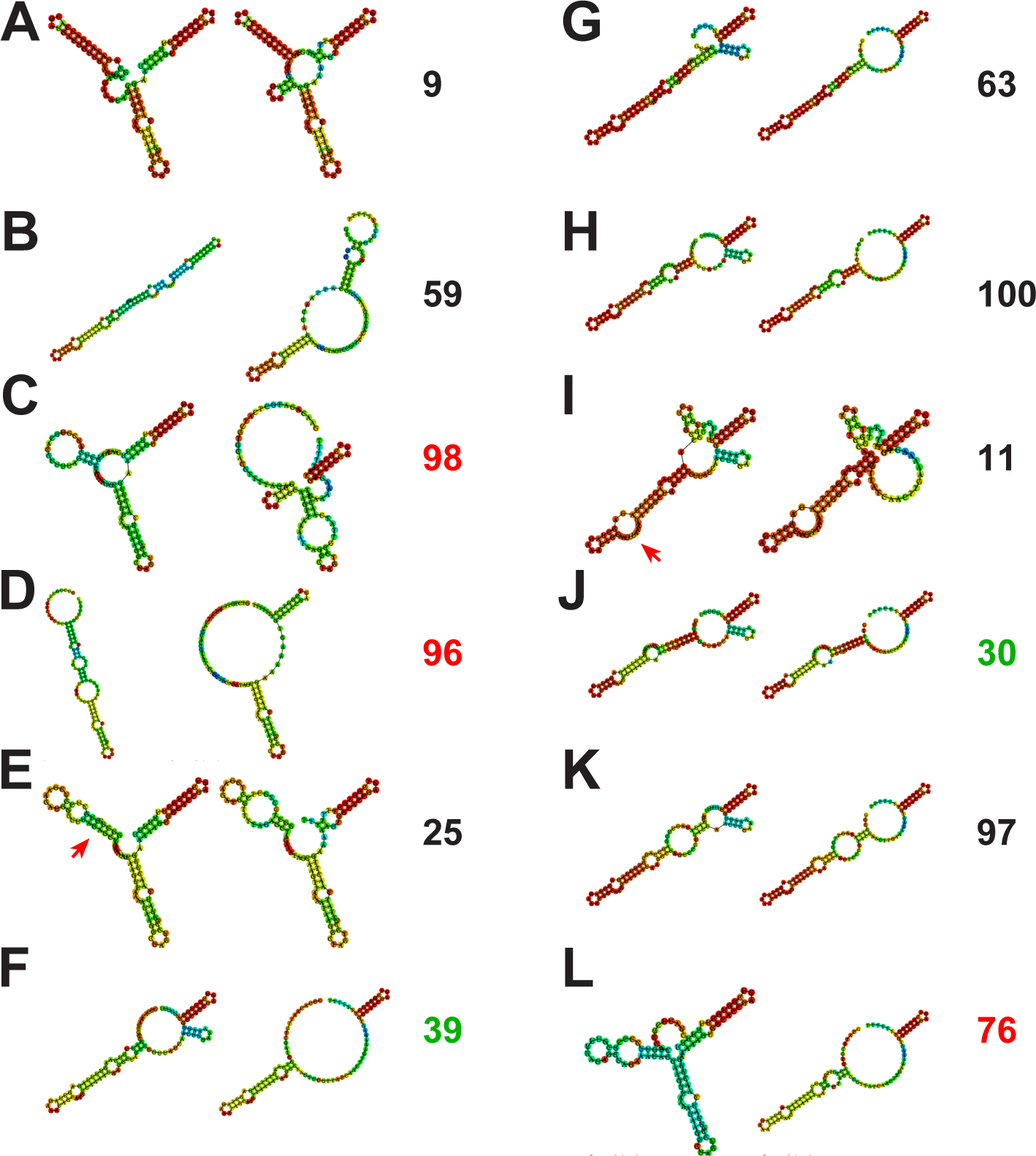
Secondary structure of sgRNAs targeted to the Drosophila vermillion, ebony and yellow gene from Ren et al. (2014). **(A-L)** Secondary structures of sgRNAs predicted by RNAfold targeting the vermillion (A-D), the ebony (E-H) and the yellow gene (I-L). Minimal energy and proximity structures are shown on the left and right, respectively. Red and blue indicated high and low probabilities for the adopted structural base-pairing assignment, respectively. The number on the right indicates the effectiveness of inducing heritable mutations after injection into fly embryos as determined by Ren et al. (2014). Red and green numbers indicate an effectiveness which is too high or too low compared to predicted DNA cleavage efficiency. Red arrows point towards structural features likely limiting DNA cleavage.

**Supplemental Figure S5.**
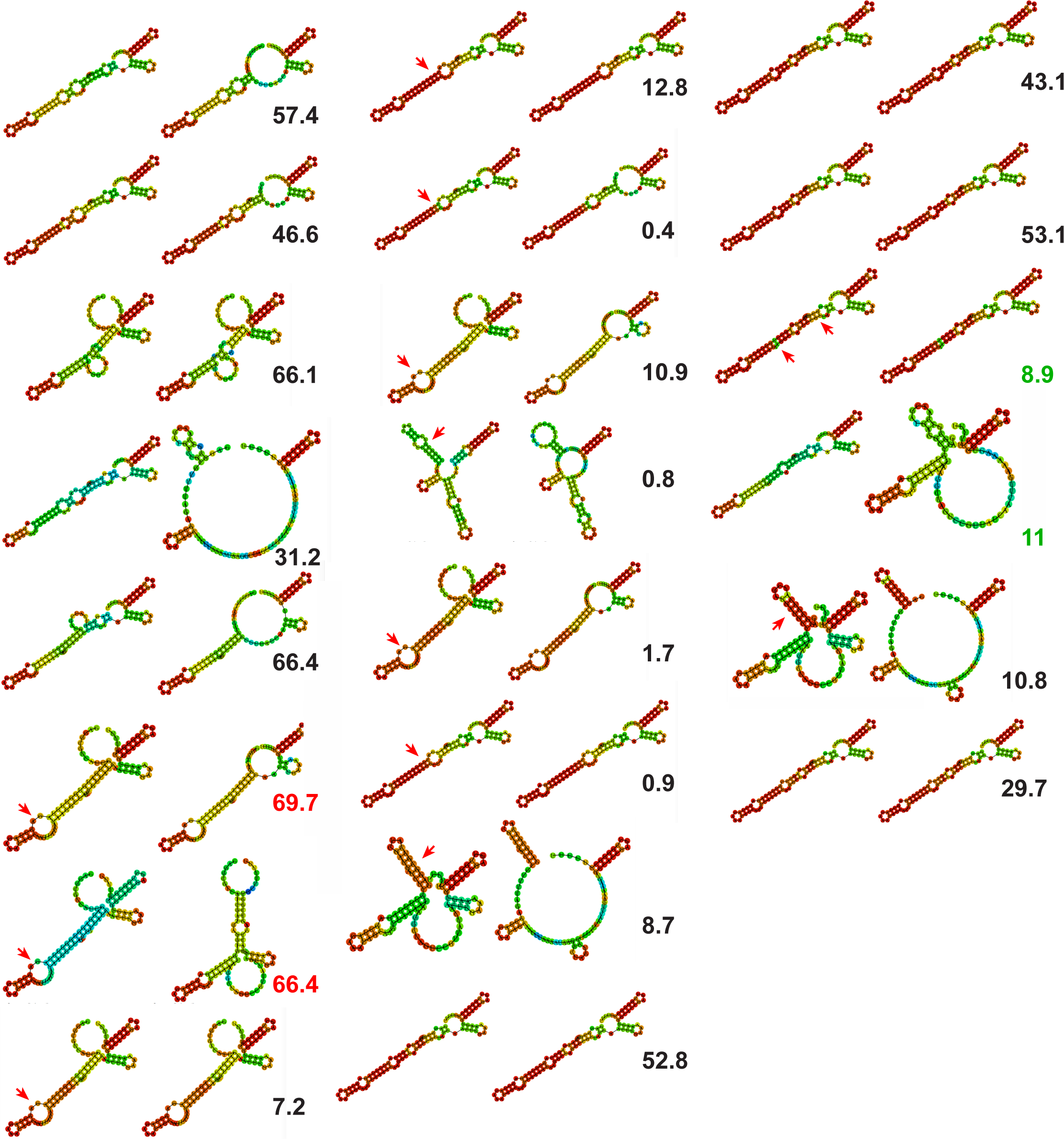
Secondary structure of sgRNAs targeted to human CD22 from Graf et al. (2019). **(A-V)** Secondary structures of sgRNAs predicted by RNAfold. Minimal energy and proximity base-pair structures are shown on the left and right, respectively. Red and blue indicated high and low probabilities for the adopted structural base-pairing assignment, respectively. The number on the right indicates the effectiveness of inducing heritable mutations after injection into fly embryos as determined by Ren et al. (2014). Red and green numbers indicate an effectiveness which is too high or too low compared to predicted DNA cleavage efficiency. Red arrows point towards structural features likely limiting DNA cleavage.

**Supplemental Figure S6.**
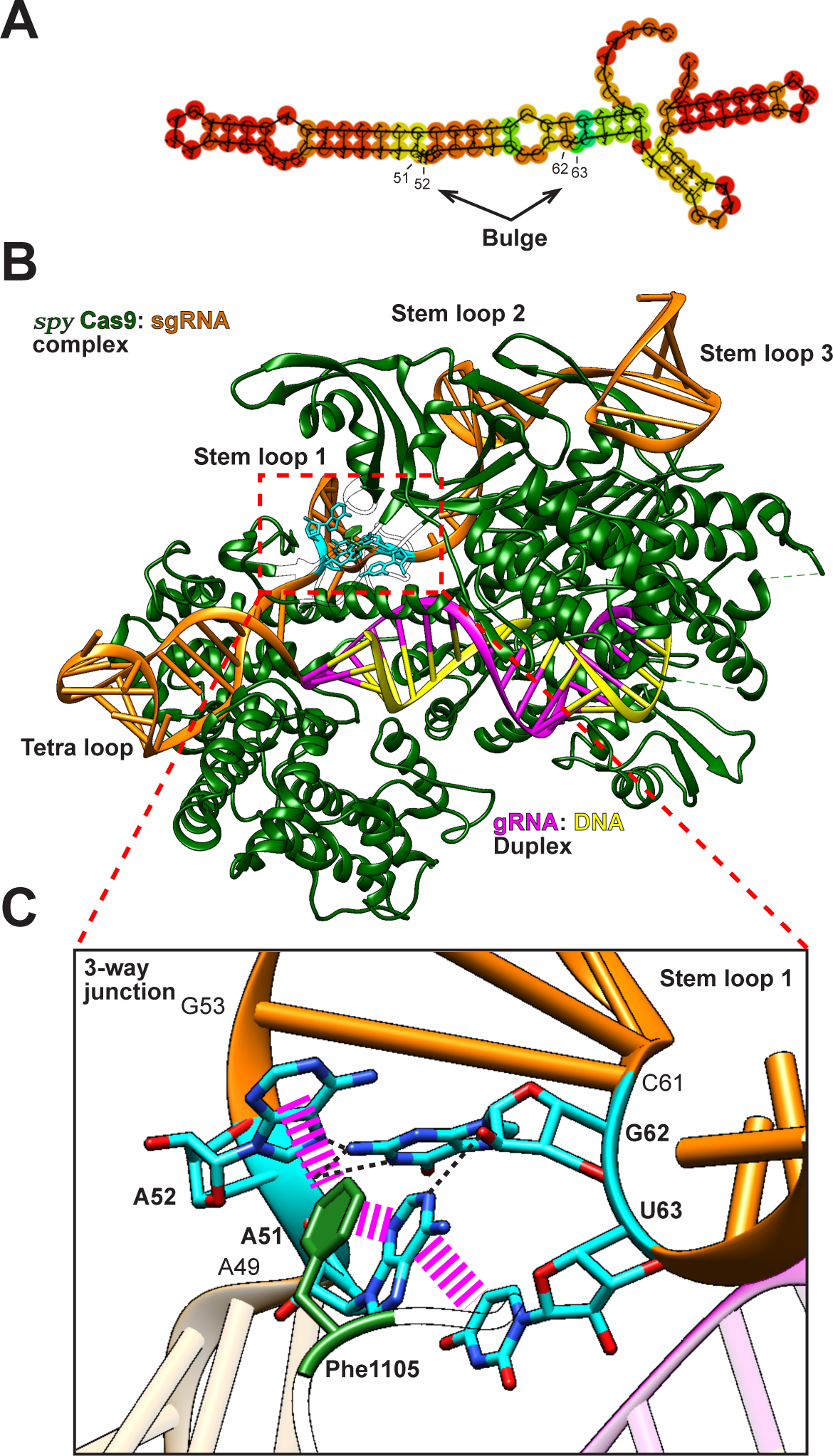
Key sgRNA secondary structural features directly interact with the Spy Cas9 endonuclease in the three way junction. **(A)** Minimum Free Energy (MFE) predicted secondary structure of the sgRNA used for co-crystallization of Spy Cas9 (Nishimasu et al., 2014). Nucleotides involved in the three way junction interactions are indicated (A_51_, A_52_, G_62_ and U_63_). **(B)** X-ray crystal structure of†spyCas9 endonuclease in complex with chimeric sgRNA bound to genomic DNA target (PDB: 4OO8) (Nishimasu et al 2014). The gRNA portion is coloured in pink and the remainder of the sgRNA is coloured in orange.Genomic DNA is coloured yellow and the protein chain is coloured in green. Nucleotide residues situated at key structural features in the sgRNA are coloured cyan. Note that a small portion of the Cas9 protein chain depicting amino acid side chains to view the sgRNA three-way junction is transparent.† **(C)** Magnified view of the sgRNA three-way junction. Hydrogen bonds are indicated by black dashed lines and aromatic stacking interactions are shown by pink dashed lines. Here, the interaction of Phe_1105_ with A_51_, A_52_, and U_63_ and the interaction of G_62_ with A_51_, A_52_ and and the phosphate backbone can be seen.

**Supplemental Figure S7.**
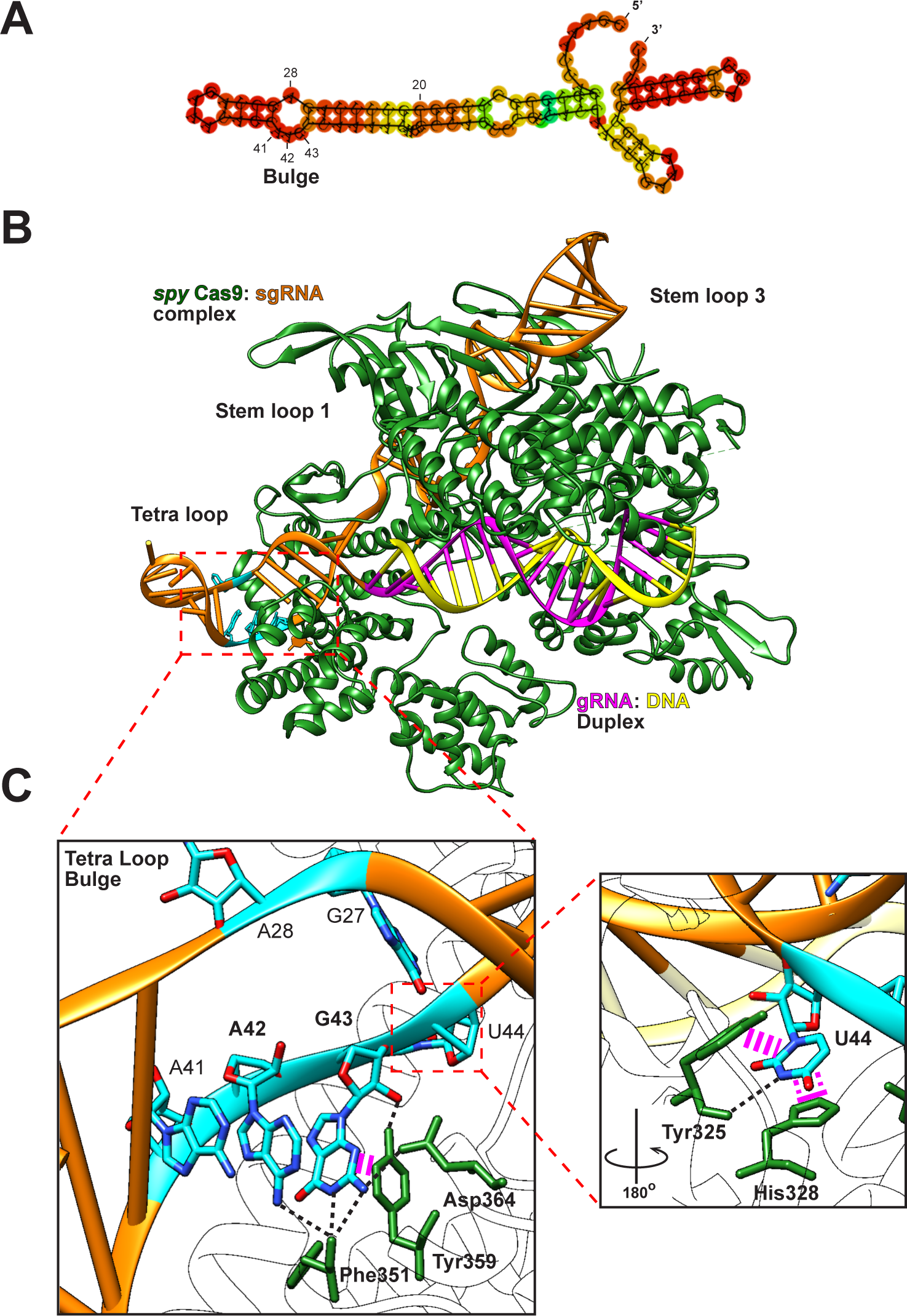
Key sgRNA secondary structural features directly interact with the Spy Cas9 endonuclease in the tetraloop bulge. **(A)** Minimum Free Energy (MFE) predicted secondary structure of the sgRNA used for co-crystallization of spyCas9 (Nishimasu et al., 2014). Nucleotides forming the bulge are indicated (A_28_, A_41_, A_42_ and G_43_). **(B)** X-ray crystal structure of Spy Cas9 endonuclease in complex with chimeric sgRNA bound to genomic DNA target (PDB: 4OO8) (Nishimasu et al 2014). The gRNA portion is coloured in pink and the remainder of the sgRNA is coloured in orange. Genomic DNA is coloured yellow and the protein chain is coloured in green. Nucleotide residues situated at key structural features in the sgRNA are coloured cyan. Note that the tetra loop after the bulge does not interact with Cas9 and sticks out of the structure. **(C)** Magnified view of the sgRNA bulge present in the tetra loop. Hydrogen bonds are indicated by black dashed lines and aromatic stacking interactions are shown by pink dashed lines. Here, the interactions of Phe_351_, Tyr_359_ and Asp_364_ with A_42_ and G_43_ can be seen. Inlet on the right: 180° turn to visualize base-stacking of U_44_ with Tyr_325_ and His_328_. Through these interactions base-pairing with G_27_ is prevented.

**Supplemental Figure S8.**
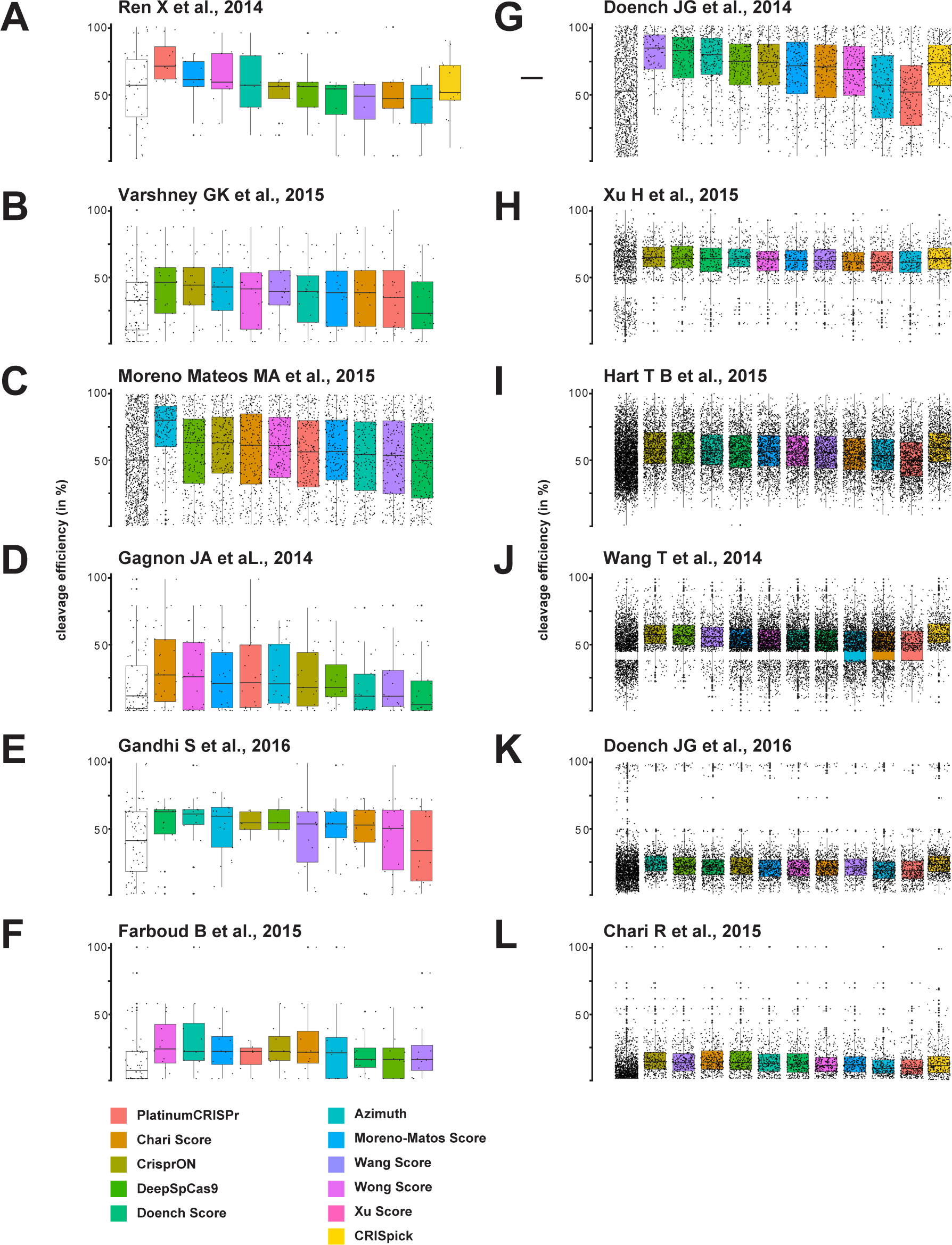
Comparison of sgRNA selection by different sgRNA selection tools shown as median of the cleavage efficiency for individual data sets (A-L). The distribution of cleavage efficiencies for all sgRNAs is shown on the left (white box).

**Supplemental Figure S9.**
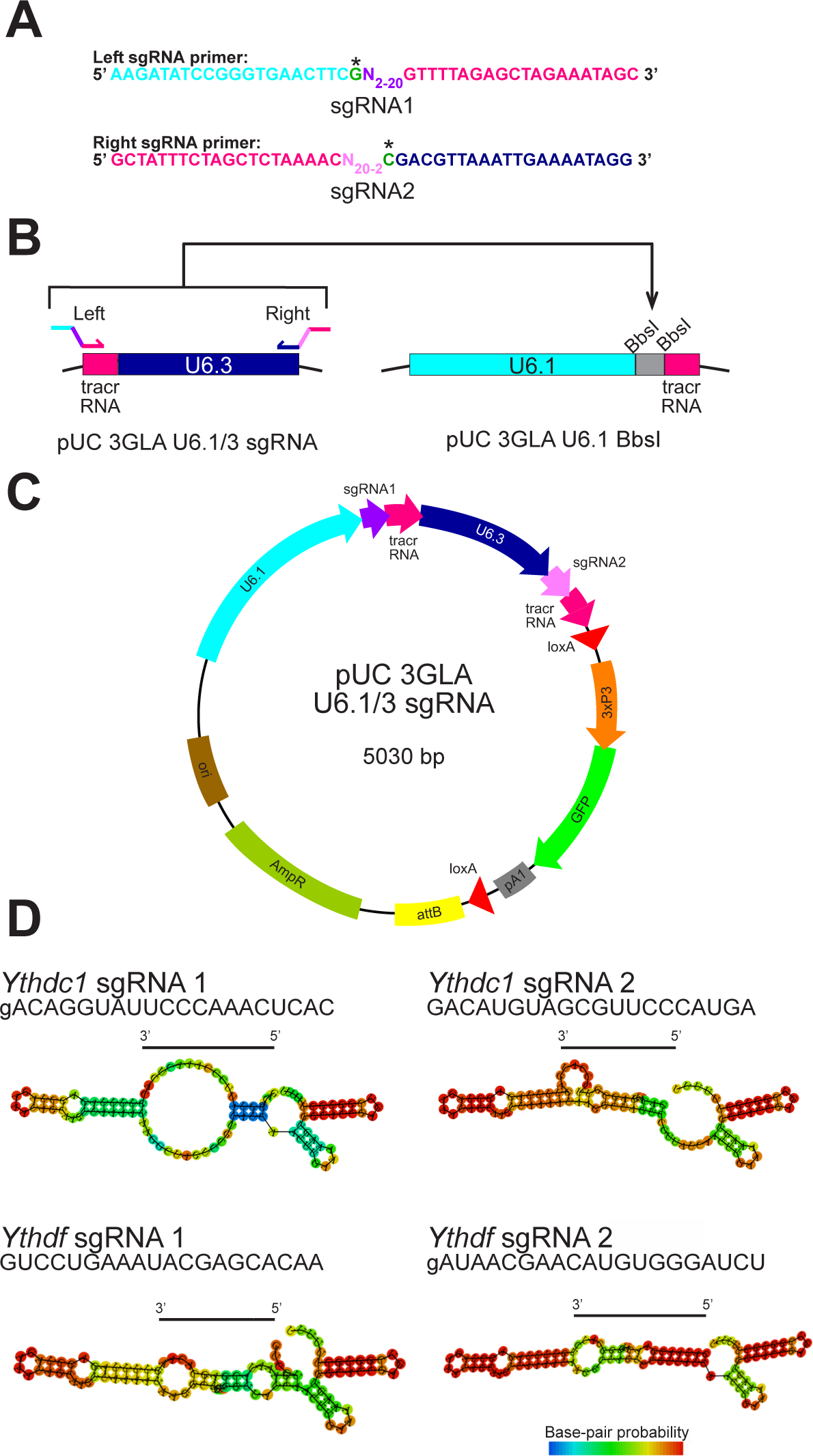
*Drosophila* GFP-marked transformation vector for U6 promoter mediated expression of two sgRNAs. **(A)** Forward and return primer sequences to incorporate sgRNA sequences. The first nucleotide of the gRNA is indicated by an asterisk. **(B)** Cloning scheme for incorporating two sgRNAs into the destination vector. **(C)** Plasmid map of the fly transformation vector *pUC 3GLA U6.1/U6.3* sgRNA expressing two sgRNAs under U6.1 and U6.3 promoters, respectively. **(D)** Secondary structure of sgRNAs targeting *Ythdc1* and *Ythdf*.

**Supplemental Figure S10.**
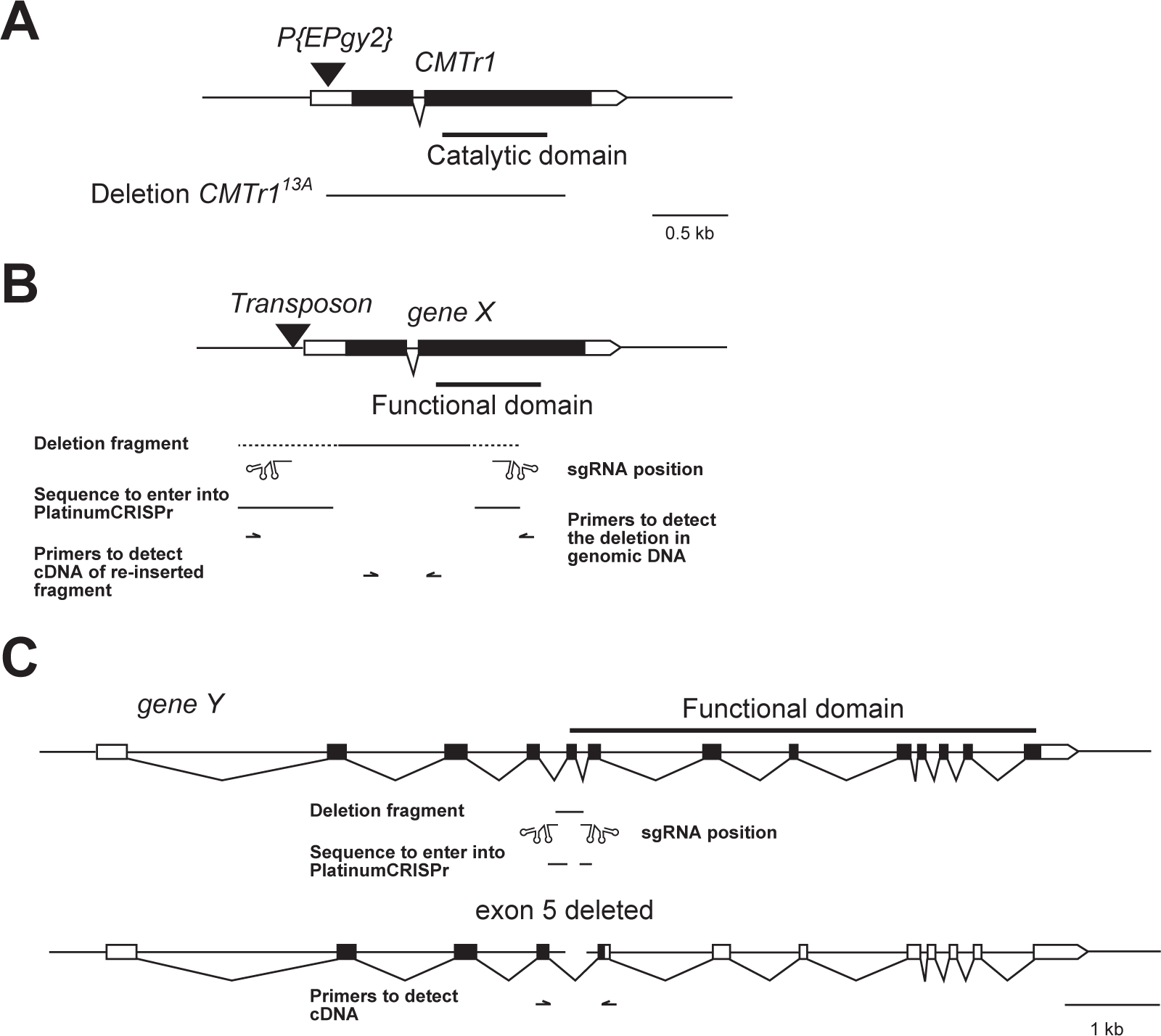
Generation of gene knock-outs by targeted deletions. **(A)** Schematic of a the *Drosophila CMTR1* gene indicating transcribed parts in boxes with translated parts in black. Gaps between boxes indicate introns which are spliced out. The triangle indicates a positively marked transposon. The *w+* marked P element was mobilized over a *w+* marked chromosomal deficiency and a deletion starting at the P element insertion site and extending into the catalytic domain was recovered (deletion *CMTr1*^13A^). **(B)** Schematic of a typical *Drosophila* gene indicating the position of sgRNAs and a deletion to be generated to obtain a knock-out below the gene model. The line indicates the desired deletion and flanking dashed lines indicate positions where gRNAs can be selected. Below are primers indicated to screen for the deletion which will result in a short PCR product if the deletion is present compared to a long PCR product in the absence of a deletion, which will likely not amplify efficiently. Then, to screen for re-insertions, primers are indicated flanking an intron in the deleted part. If re-integration occurs, a short PCR product will be amplified for integration in the sense direction. If a longer PCR product is amplified, indicating a non-spliced intron, integration will be in the anti-sense direction. **(C)** Schematic of a typical human gene composed of short exons and large introns with the translated part marked in black. Removal of an exon can lead to a frameshift resulting in a truncated protein, that is non-functional (below). The line indicates the desired deletion induced by the indicated sgRNAs. RT-PCR with primers in exons flanking the deleted exon can be used to monitor CRISPR/Cas9 induced deletion efficiency in cultured cells based on the ration of splice products.

## Notes

### Competing Interest Statement

The authors have declared no competing interest.

### Summary of Updates

Additional data and analysis was added.

